# A Kink in DWORF Helical Structure Controls the Activation of the Sarco-plasmic Reticulum Ca^2+^-ATPase

**DOI:** 10.1101/2021.05.05.442831

**Authors:** U. Venkateswara Reddy, Daniel K. Weber, Songlin Wang, Erik K. Larsen, Tata Gopinath, Alfonso De Simone, Seth Robia, Gianluigi Veglia

**Author notes:** To whom correspondence should be addressed: Gianluigi Veglia, Department of Biochemistry, Molecular Biology & Biophysics, University of Minnesota, 6-155 Jackson Hall, MN 55455.

## Abstract

The sarco(endo)plasmic reticulum Ca^2+^-ATPase (SERCA) is a P-type ATPase embedded in the sarcoplasmic reticulum. For each enzymatic cycle, SERCA transports 2 Ca^2+^ ions per ATP hydrolyzed in exchange for 2 to 3 H^+^ ions. SERCA is responsible for approximately 70% of Ca^2+^ transport and plays a central role in muscle relaxation. SERCA’s function is regulated by endogenous regulins, single-pass membrane proteins that bind the ATPase within the membrane. While most of the regulins, such as phospholamban and sarcolipin, inhibit SERCA’s activity, a newly discovered protein DWarf Open Reading Frame (DWORF) has a unique activating effect. DWORF is a 3.8 kDa bitopic membrane protein expressed in cardiac muscle. In this work, we determine the structure, topology, and per-residue lipid interactions of DWORF in lipid bilayers using a combination of high-resolution oriented sample solid-state NMR (OS-ssNMR) spectroscopy and refinement by replica-averaged orientationally-restrained molecular dynamics (RAOR-MD). We found that DWORF’s structural topology consists of a dynamic N-terminal domain, an amphipathic juxtamembrane helix that crosses the lipid groups at an angle of 64° and a transmembrane (TM) C-terminal helix with an angle of 32°. A kink induced by Pro15, unique to DWORF, separated the two helical domains. A single Pro15Ala mutant significantly decreases the kink and eliminates DWORF’s activating effect on SERCA. Overall, our findings directly link DWORF’s structural topology to its unique activating effect on SERCA.

## Introduction

Cardiac muscle contraction is controlled by an array of Ca^2+^-handling proteins that regulate dynamic changes in cytoplasmic Ca^2+^. In myocytes, contraction is initiated by the entry of extracellular Ca^2+^ during an action potential, which in turn triggers the ryanodine receptors (RyR) to release Ca^2+^ stored in the sarcoplasmic reticulum (SR) in a process referred to as calcium-induced calcium release (CICR). Muscle relaxation occurs when Ca^2+^ levels fall active transport of Ca^2+^ back into the SR by the sarco(endo)plasmic reticulum Ca^2+^-ATPase (SERCA) at the expense of ATP hydrolysis, which is responsible for 70% Ca^2+^ removal (Luo and Anderson, 2013, Bers, 2002). SERCA is a 110 kDa P-type ATPase that pumps Ca^2+^ ions by oscillating between a high-Ca^2+^-affinity (*E1*; open to cytoplasm) and low-Ca^2+^ (*E2*; open to SR lumen) conformational states. SERCA is co-expressed with small bitopic mini-membrane proteins (< 100 amino acids) that bind within a regulatory intramembrane groove positioned between the transmembrane (TM) domains 2, 6, and 9 of SERCA, which fine-tunes its activity in a tissue-specific manner (Makarewich, 2020, Zhihao et al., 2020).

In skeletal muscle, the SERCA1a isoform is regulated by sarcolipin (SLN). This 31amino acid polypeptide has been linked to the uncoupling of ATP hydrolysis from Ca^2+^ transport of SERCA, contributing non-shivering thermogenesis and whole-body metabolism (Periasamy et al., 2017, Palmer and Clegg, 2017, Gamu et al., 2020). On the other hand, the SERCA2a isoform in cardiac muscle is co-expressed with phospholamban (PLN), a 52 amino acid membrane protein. PLN is one of the key targets of the β-adrenergic pathway responsible for the fight-of-flight response. Phosphorylation of PLN relieves its inhibition of SERCA, increasing Ca^2+^ transport during exercise or other physiological stress (Kranias and Bers, 2007, MacLennan and Kranias, 2003, Kranias and Hajjar, 2017). SLN has also been identified in the atria and has been linked to cardiac dysfunction (Morales Rodriguez et al., 2020, Shanmugam et al., 2015, Zheng et al., 2014), although its functional role in the heart is not entirely clear. Recently, other SERCA-binding regulins have been discovered, and this family now includes myoregulin (MLN; co-expressed with SERCA1a in skeletal muscle), another regulin (ALN; co-expressed ubiquitously with SERCA2b), endoregulin (ELN; co-expressed with SERCA3 in endothelial and epithelial cells), and dwarf open reading frame (DWORF; co-expressed with SERCA2a and PLN in cardiac muscle) (Anderson et al., 2016, Nelson et al., 2016, Singh et al., 2019, Anderson et al., 2015).

While all regulins generally inhibit SERCA by decreasing its apparent Ca^2+^-binding affinity, DWORF is the only one found to be deemed an ‘activator’ (Nelson et al., 2016). DWORF’s activation is hypothesized to occur by outcompeting the binding of PLN to SERCA, essentially reviving SERCA’s uninhibited Ca^2+^-binding affinity (Nelson et al., 2016, Makarewich et al., 2018). This was suggested by cardiomyocytes obtained from DWORF-overexpressing transgenic (Tg) mice having a similar phenotype to those obtained from PLN-null mice, such as faster cytosolic Ca^2+^ decay rates, and a higher Ca^2+^ SR load (Makarewich et al., 2018, Nelson et al., 2016). Regulator displacement has also been demonstrated by FRET and co-immunoprecipitation studies using cell lines co-expressing SERCA, PLN, SLN, MLN and DWORF (Makarewich et al., 2018, Nelson et al., 2016, Li et al., 2021). DWORF mRNA levels are also reduced in human ischemic heart failure and in mouse models of cardiovascular disease (Nelson 2016). Evidence of direct activation of SERCA by DWORF (*i*.*e*., in the absence of PLN) has also been recently observed (Li et al., 2021). Since the augmentation of Ca^2+^ transport in the SR constitutes a general strategy for improving cardiac contractility in heart disease patients, DWORF represents a potential target to ameliorate muscle disease. Indeed, DWORF overexpression in a mouse model for dilated cardiomyopathy ameliorates diastolic dysfunction and reverses the progression of heart failure (Makarewich et al., 2018). These studies have generated intense interest in the structure/function mechanisms governing SERCA regulation by DWORF. High-resolution structural data are needed to understand fundamental determinants of DWORF’s unique regulatory activity and create a basis for rational development of DWORF as a potential treatment for heart failure.

Here, we utilized a combination of oriented sample solid-state NMR (OS-ssNMR) and dynamic refinement in explicit membranes to determine the structure and membrane architecture of DWORF in lipid bilayers. We found that this small membrane protein adopts a helical conformation similar to other regulins, but with a distinctive proline-induced (Pro15) kink separating the amphipathic N-terminal end of the helix from the C-terminal transmembrane (TM) portion. By mutating Pro15 into alanine, DWORF adopts a continuous helix, with a less pronounced kink. Remarkably, this single mutation abolishes DWORF’s activating effect on SERCA’s maximum enzymatic turnover rate (V_max_). These structural features may account for the functional differences that distinguish DWORF from the other regulators.

## RESULTS

### Structural topology of DWORF in lipid membranes as determined by MAS and OS-ssNMR spectroscopy

DWORF was overexpressed in *E. coli* bacteria as a soluble fusion with maltose-binding protein (MBP) and purified as we have reported previously for both PLN and SLN (Buck et al., 2003) (see **Figure 1A** and **Figure S1** for sequence, gel and MALDI analysis). For OS-ssNMR samples, DWORF was reconstituted into lipid bicelles composed of DMPC (1,2-dimyristoyl-sn-glycero-3-phosphocholine), POPC (1-pal-mitoyl-2-oleoyl-sn-glycero-3-phosphocholine), and PE-DTPA (1,2-dimyristoyl-sn-glycero-3-phosphoethanolamine-N-diethylenetriaminepentaacetic acid) at a molar ratio of 9.8:2.7:0.2. To form the bicellar phase DHPC (1,2-dihexanoyl-sn-glycero-3-phosphocholine) was added to the lipid mixture, q = 4.4. These anisotropic bicelles with reconstituted proteins align relative to the direction of the static magnetic field, and the anisotropic NMR parameters become orientationally dependent. Solid-state separated local fields (SLF) experiments enable one to measure ^15^N chemical shift anisotropy (CSA) and ^1^H-^15^N dipolar couplings (DC) for each amide group. The CSA and DC values can be used directly to calculate the structure and topology (*i*.*e*., tilt and azimuthal angles) for helical and β-sheet membrane proteins (Wang et al., 2000, Ramamoorthy et al., 2004). The sensitivity-enhanced (SE)-SAMPI4 SLF experiment (Gopinath et al., 2010b) was used to image the oriented amide fingerprint of DWORF (**Figure 1B**). Higher resolution spectra were obtained using bicelle-protein complexes *flipped* with the normal of the membrane parallel to the direction of the static magnetic field by chelating Yb^3+^ ions with PE-DTPA lipids (Prosser et al., 1998). This expedient effectively doubles the spectral resolution compared to *unflipped* samples (**Figure S2**). The uniform orientation of the bicelle sample was confirmed by ^31^P NMR, where the more intense peak of the DMPC lipid head groups, showing a rather uniform alignment of the membrane bilayer (**Figure S2**). (SE)-SAMPI4 spectra resulted in well-resolved resonances displaying periodic patterns (PISA wheels) of CS and DC typical of TM proteins with regular α-helical structure (Ramamoorthy et al., 2004). Spectral resonances were assigned using twelve selectively labeled samples (**Figure S3**). Using these resonances as a seed, we carried out unrestrained MD simulations of truncated DWORF (DWORF_7-35_) and modeled as an ideal α-helix. After a short equilibration, we back-calculated the SLF spectra and used it to guide the resonance assignments (**Figure S4**). The explicit membrane environment used in the simulations drives the topology of DWORF to form two distinct helical segments kinked at Pro15. The predicted and experimental CSA and DC are in good agreement, with several resonances located in the C- and N-terminal helical segments overlaying almost perfectly. To complete the structural characterization, we carried out magic angle spinning (MAS) experiments on U-^13^C,^15^N DWORF recon-stituted into *d*_54_-DMPC liposomes. The cross-polarization (CP)-based SLF experiments detect the most rigid regions of the protein. However, the [^13^C-^13^C] DARR and [^13^C-^15^N] NCA MAS spectra display the limited resolution of helical membrane proteins (**Figure S5**), hampering sequential assignments. Nonetheless, the 2D [^13^C-^13^C] TOBSY experiment was able to detect the dynamic N- and C-terminal residues (**Figure 1C**). When combining both OS-ssNMR and MAS data we identified three distinct domains: a dynamic N-terminus (Ia; Met1-Ala5), an amphipathic juxtamembrane domain (Ib; Gly6 to Leu12), and a hydrophobic TM domain (II, Leu13 to Ser35). Using the structural fitting of the NMR resonances (PISA wheel), we determined the topology of DWORF, whose helical domain Ib adopts a tilt (θ_Ib_) and azimuthal angles (ρ_Ib, F9_; referenced to Phe9) of 63.6 ± 2.0° and 129 ± 17°, respectively. The fitting of domain II resonances in the downfield region of the spectrum (130 to 200 ppm; **Figure 1B**) show that this domain assumes a tilt (θ_II_) and azimuthal (ρ_Ib, V24_) angles of 33.1 ± 0.5° and 105 ± 4°, respectively.

**Figure 1.**
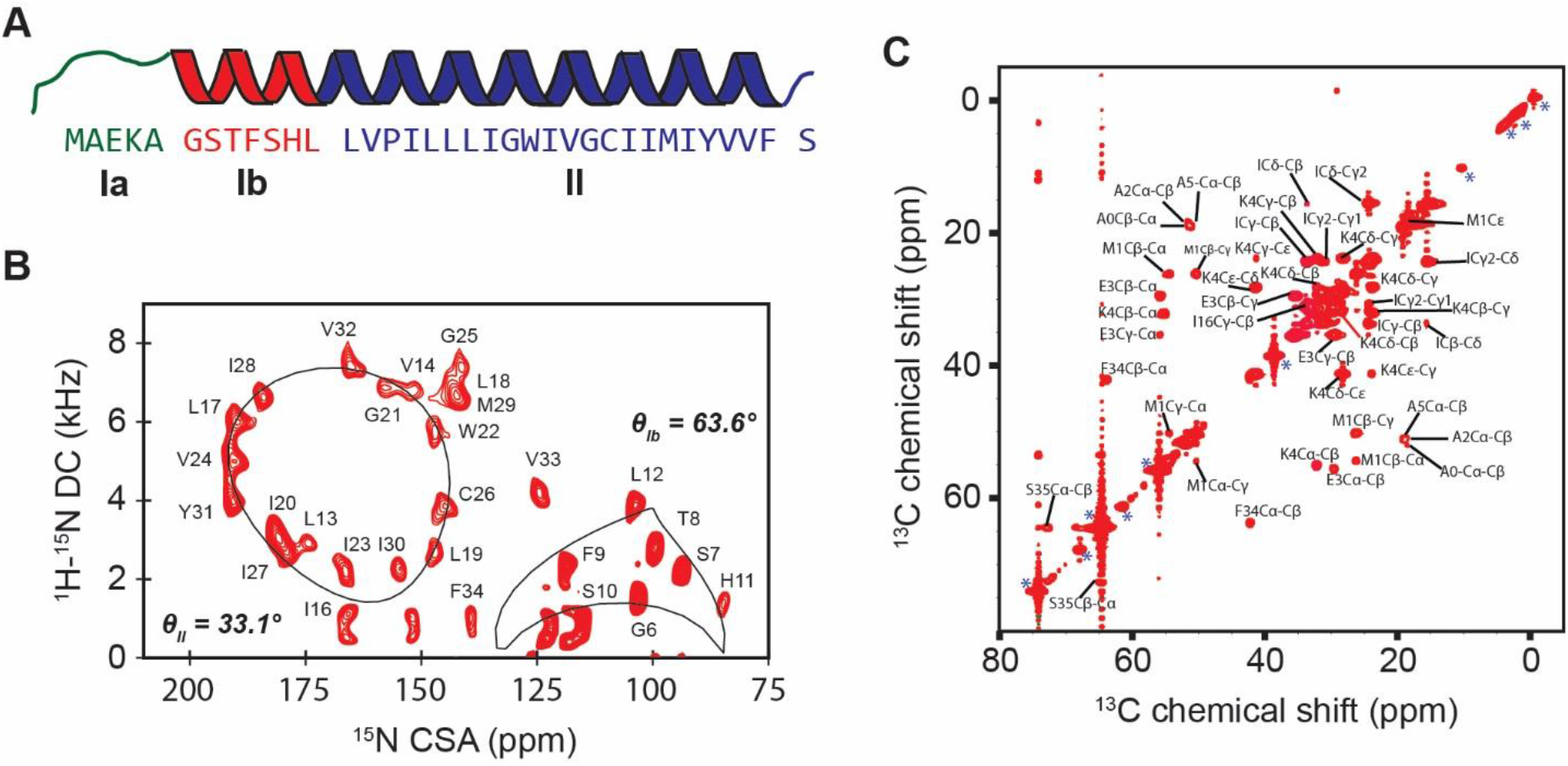
**(A)** Amino acid sequence, secondary structure and domain nomenclature of DWORF. **(B)** Assigned (SE)-SAMPI4 OS-ssNMR spectrum of ^15^N-DWORF reconstituted into *flipped* bicelles. PISA-wheel fits (lines) and tilt angles for domains Ib (θ_Ib_) and II (θ_Ib_) are shown. **(C)** ^13^C TOBSY MAS spectrum showing dynamic residue of ^13^C,^15^N-DWORF reconstituted into DMPC liposomes.

### Calculations of the structure and topology of DWORF using anisotropic NMR parameters

Assigned ^15^N CSA and ^1^H-^15^N DC correlations in SLF spectra (summarized in **Table S1**) related directly to the orientation of backbone amides and were used as the primary source of topological information for simulated annealing structure calculations (Shi et al., 2009, Weber et al., 2020a). Close agreement of CS and DC to the PISA model allowed us to assume an α-helical structure from residues Gly6 to Phe34, which were enforced by idealized torsion angles with generous boundaries to allow for structural perturbations. An empirical membrane potential with a hydrophobic thickness equivalent to the bicelle lipid composition was applied after simulated annealing stages to obtain the correct depth of insertion (Senes et al., 2007). **Figure 2A** shows the refined conformational ensemble, which converged to the same membrane insertion depth. All conformers display a pronounced kink separating domains Ib and II. Tilt angles were 63.8 ± 1.6° and 31.9 ± 1.1° for domains Ib and II, respectively, and their corresponding azimuthal angles of 129.7 ± 3.0° and 95.8 ± 2.2° at Phe9 and Val24, respectively, matching the structural fitting obtained with the PISA wheel method (Marassi and Opella, 2000, Murray et al., 2014). The amphipathic domain Ib (Gly6 to Leu12) is adsorbed onto the lipid headgroup, with Phe9 acting as an anchor to the membrane. Domain II (Leu13 to Ser35) transverses the acyl layer with a fixed rotation. The C-terminal Tyr31 and Ser35 *snorkel* at the membrane-water interface. While our data cannot confirm DWORF oligomerization, fluorescence resonance energy transfer (FRET) studies suggest the presence of dimers or higher order oligomerization (Singh et al., 2019). According to our calculated structural ensemble, the putative oligomerization sequence (G_21_XXXG_25_) is located in the middle of domain II, causing a slight curvature, which may also be prone to dimerization (Singh et al., 2019). The complete statistics of the final structural ensemble are summarized in **Table 1**. The calculations converged to an average backbone rmsd of 0.3 Å (if domain Ib and II are fitted separately), while the side-chains were less defined (rmsd 1.9 Å and 2.1 Å for domains Ib and II, respectively) as the only restraints included in the calculations were imposed by the covalent geometry and knowledge-based torsion angle potentials (Bermejo et al., 2012). The experimental CSA and DC values were reproduced with *R*-factors (Kim et al., 2001) of 0.59 and 0.73, respectively (**Figure 2B, Table 1**).

**Figure 2.**
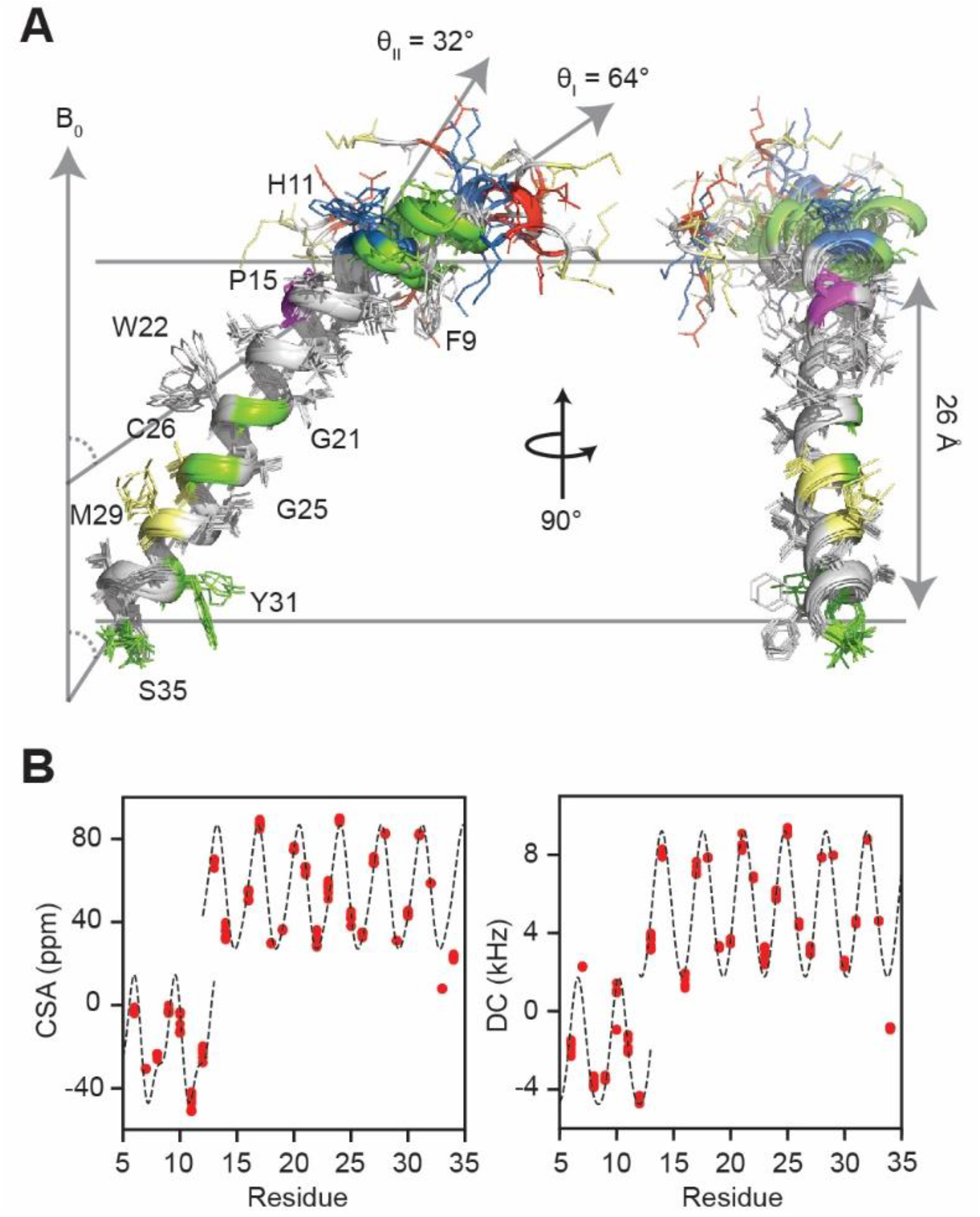
**(A)** Overlay of the 10 lowest energy structures of DWORF. Structures were aligned according to the center of mass of residues 13 to 35 without translations or rotations affecting the depth-of-insertion, tilt (θ) or azimuthal angle (ρ). Calculations were performed using XPLOR-NIH with a 26 Å virtual membrane equivalent to the hydrophobic thickness of the experimental DMPC/POPC bicelle composition. Polar (green), aromatic, basic (blue), proline (magenta), acidic (red) and sulfur-containing (yellow) residues are labeled. (**B)** ^15^N CSA and ^15^N-^1^H DC wave plots of back-calculated values from the 10 structural models. Broken lines mark the PISA wheel fits to experimental restraints scaled to account for a general order parameter of 0.8. CSA values are shown in the reduced form subtracting the isotropic chemical shift.

**Table 1:** Summary statistics of 10 lowest energy structures.

| Protein Geometry * |  |
| --- | --- |
| Poor rotamers, % | 5.7 ± 4.7 |
| Favored rotamers, % | 89.7 ± 5.3 |
| Ramachandran outliers, % | 0.6 ± 1.3 |
| Ramachandran favored, % | 96.1 ± 3.2 |
| RMSD (heavy atoms), Å † |  |
| All (Res. 6-34) |  |
| Backbone | 3.07 ± 0.53 |
| Side-chain | 2.45 ± 0.35 |
| Domain I (Res. 6-12) |  |
| Backbone | 0.26 ± 0.09 |
| Side-chain | 1.93 ± 0.34 |
| Domain II (Res. 13-34) |  |
| Backbone | 0.33 ± 0.13 |
| Side-chain | 2.12 ± 0.40 |
| Restraints |  |
| Dihedral angles (ϕ, ψ) ‡ | 56 |
| Hydrogen bonds ‡ | 21 |
| CSA | 28 |
| DC | 28 |
| CSA R-factor | 0.59 ± 0.05 |
| DC R-factor | 0.73 ± 0.05 |
| Protein Topology § |  |
| Domain I θ, ° | 63.8 ± 1.6 |
| Domain I ρ, ° (F9) | 129.7 ± 3.0 |
| Domain II θ, ° | 31.9 ± 1.1 |
| Domain II ρ, ° (V24) | 95.8 ± 2.2 |

\* Determined for restrained residues Gly6 to Phe34 using Molprobit (Williams et al., 2018). † Average and standard deviations determined from fitting to the lowest energy structure. ‡ Synthetic dihedral (ϕ = -63°, ψ = -42°) were applied to helical residues Gly6 to Phe34. Dihedrals at the kink connecting Leu12 and Leu13 were unrestrained. Intradomain 1-4 N-H-O hydrogen bond restraints were also used to reinforce helical structure. § Tilt angles (θ) were determined from two center of masses (N, C and Cα atoms) of N- and C-terminal halves of the helical segment. Azimuthal angles (ρ) were determined from the coordinate of the preceding Cα atom of the respective residue to be consistent with the definition used by PISA-SPARKY (Weber et al., 2020b).

### Proline-induced kink of DWORF drives SERCA activation

To understand the functional relevance of DWORF’s kinked topology, we engineered a Pro15Ala mutant (DWORF^P15A^) and compared its activity with wild-type DWORF. As an internal control, we carried out ATPase assays in parallel with other regulins (**Figure 3A and 3B**). Under our experimental conditions, these assays show that DWORF reduces SERCA’s apparent calcium-binding affinity (pK_Ca_), although significantly less than PLN, but still more than SLN and phosphorated PLN (PLN^pS16^) (**Figure 3B**). The most striking effect is that DWORF enhances SERCA’s maximal activity (V_max_) by ∼29%, far beyond that of PLN (+ 8%), PLN^pS16^ (+ 7%), or SLN (+ 12%) (**Figure 3B**). From the assay curves (**Figure 3A**), the increase in V_max_ compensates for the pK_Ca_ shift and enhances SERCA activity at high Ca^2+^ concentration. Interestingly, the Pro15Ala mutation abolishes the stimulatory effect on V_max_, but retains the effect on pK_Ca_, makes it a net inhibitor at physiological Ca^2+^ concentrations.

**Figure 3.**
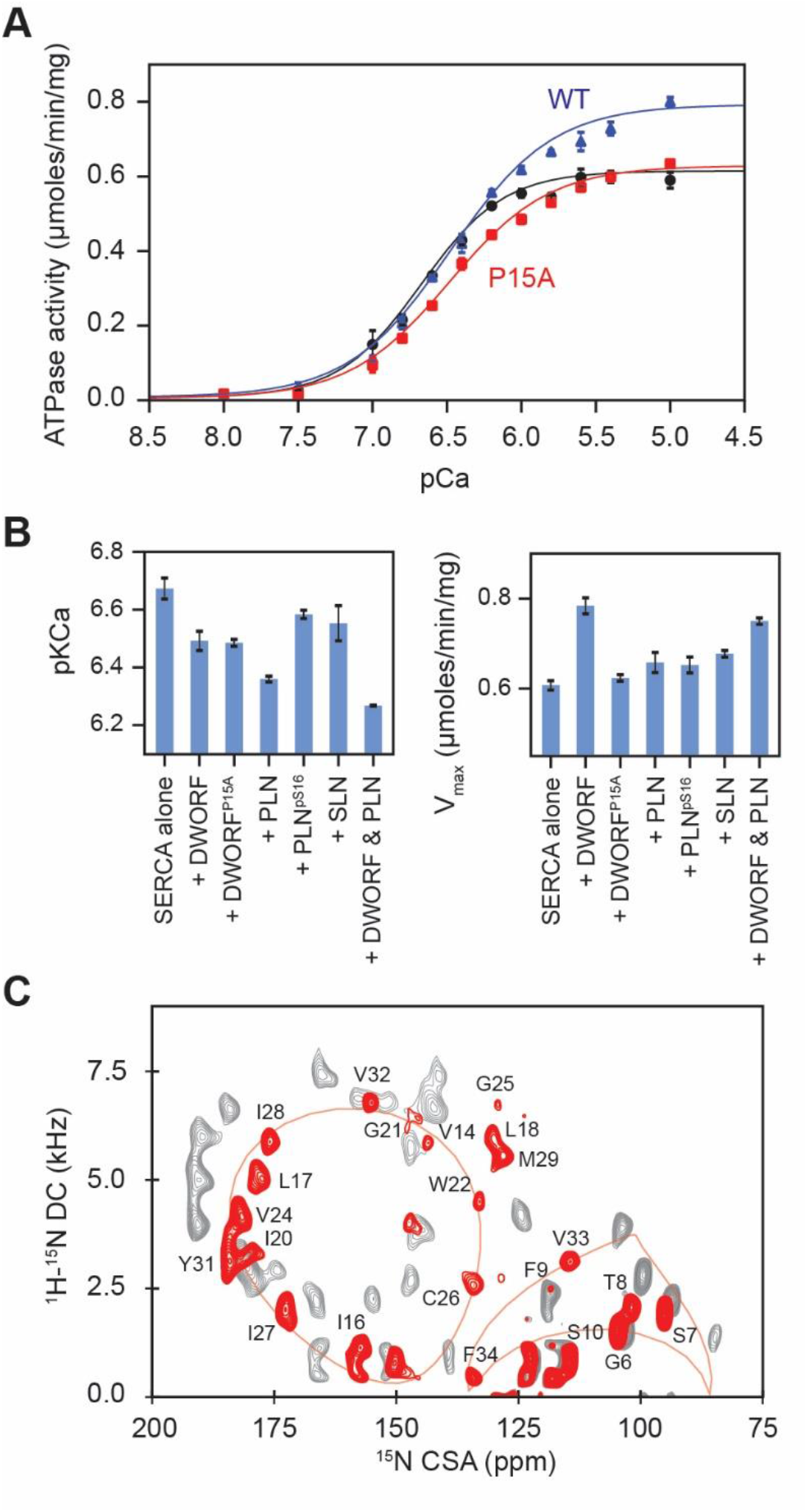
**(A)** SERCA activity alone (black circles), with DWORF (blue triangles) and DWORF^P15A^ (red squares) as function of free Ca^2+^ concentration. Curves were fitted to a Hill function to extract the Ca^2+^ concentrations at half-maximal activity (pKCa) and maximum activity (V_max_). (**B)** Summarized pKCa and V_max_ values comparing the effects of DWORF and DWORF^P15A^ to other regulators. All regulators were co-reconstituted at a 2:1 molar ratio to SERCA. **(C)** (SE)-SAMPI4 spectrum of DWORF^P15A^ (red) overlaid on wild-type DWORF (grey).

The structural analysis of DWORF^P15A^ using an (SE)-SAMPI4 experiment shows that the single P15A substitution reduces the pronounced kink, reducing the angle between domain II and domain Ib as shown by the shift of the PISA wheel (**Figure 3C**). PISA fitting found the new tilt for domain II to be 38.2 ± 0.6°, while the tilt of domain Ib was slightly reduced to 61.3 ± 2.0°. The resonances of His11 and Leu12 were broadened beyond detection, suggesting heterogeneity in the short hinge between domain II and domain Ib (**Figure 3C**). Taken with the elimination of the V_max_ effect, the structural data suggest that the kink between the helical domain is critical for DWORF’s activating effect on SERCA.

### Dynamic refinement of DWORF and DWORF^P15A^ in explicit lipid bilayers

The structures of DWORF and DWORF^P15A^ were refined using replica-averaged orientational-restrained molecular dynamics (RAOR-MD) (De Simone et al., 2014, Sanz-Hernández et al., 2016), generating conformational ensembles that capture the structural and dynamic properties that cannot be inferred from quasi-static structures in a implicit membrane. This approach captures the underlying topological fluctuations, side-chain dynamics, and specific lipid interactions that may influence regulatory activity. For these simulations, the lowest-energy structures of DWORF calculated using XPLOR-NIH (**Figure 3A**) were embedded in explicit DMPC/POPC bilayers to replicate our experimental bicelle composition. Each calculation comprised 8 parallel replicas (80 ns per replica) globally restrained using the experimental CSA and DC values from SLF experiments. The conformational ensemble of DWORF included 5,000 structures with Q-factors of 0.016 and 0.063 for back-calculated CSA and DC, respectively. These values are in excellent agreement with experimental data (**Figure 4A**). Tilt angles of domains Ib and II were 35.4 ± 7.6° and 64.7 ± 9.8°, respectively, with an inter-domain angle of 38 ± 10° (**Figure 4B**). Similarly, the dynamic refinement simulations reproduce well the experimental CSA and DC for DWORF^P15A^. The Q-factors for these calculations are 0.027 and 0.076 for CSA and DC, respectively (**Figure 4B**). The tilt angle for domain II is 42 ± 6°, while domain Ib is tilted by 59 ± 8°. Therefore, the kink between the helical domains is considerably less pronounced than in the wild type, with an inter-helical angle of 20 ± 8° (**Figure 4B and 4C**).

**Figure 4.**
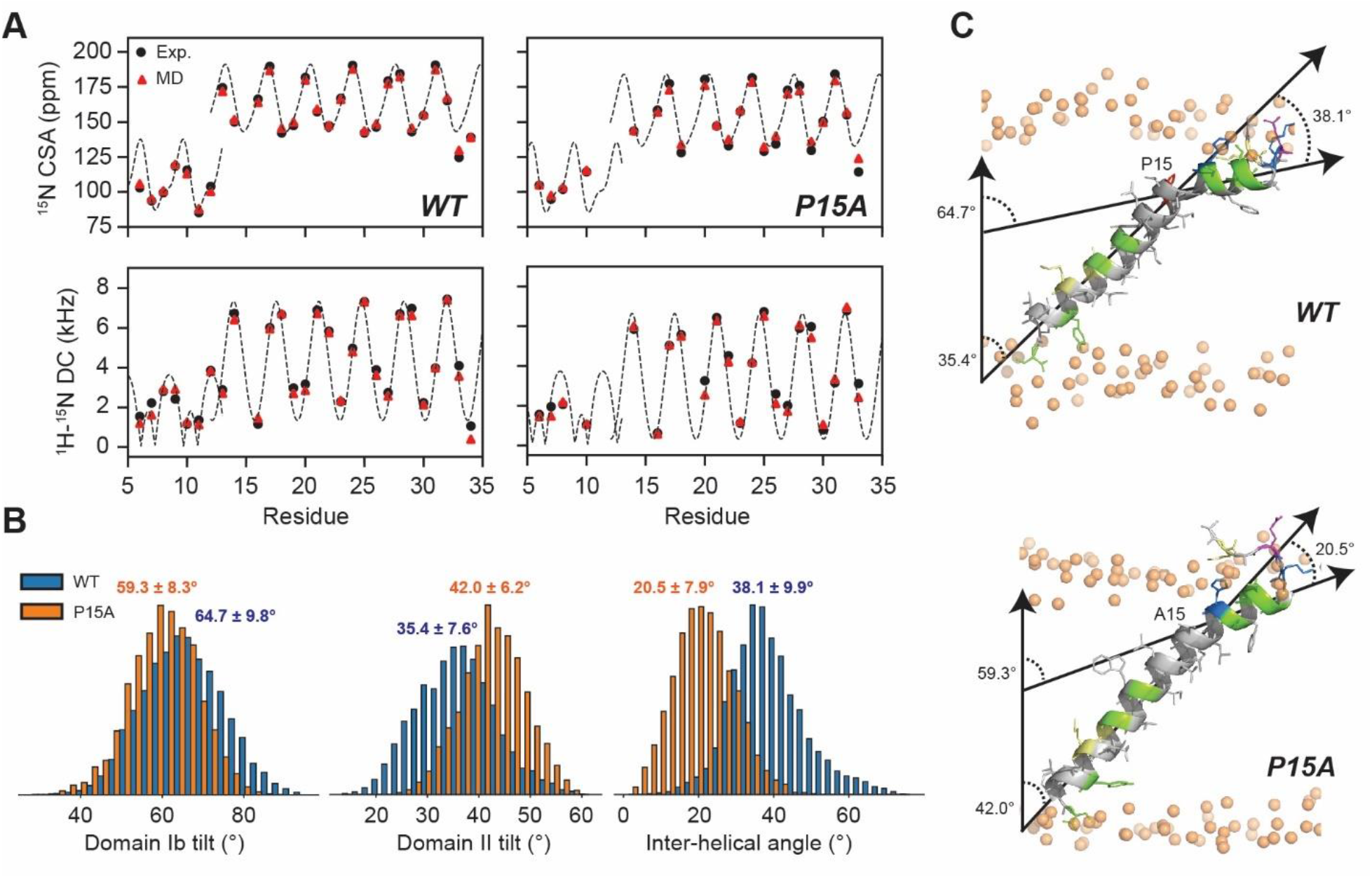
**(A)** Comparison between experimental (black) ssNMR observables and back-calculated (red) from the DWORF and DWORF^P15A^ ensembles. PISA waves fit to experimental CSA and DC are shown as broken lines. **(B)** Distributions of topological angles in DWORF (blue) and DWORF^P15A^ (orange), showing domain Ib tilt (left), domain II (middle) and inter-domain angle (right). **(C)** Representative structures of DWORF (upper) and DWORF^P15A^ (lower) displays the definitions of the topological angles.

Since the structural refinement with RAOR-MD utilizes explicit lipid bilayers, we evaluated per-residue depth-of-insertion profiles and specific lipid interactions for both DWORF^WT^ and DWORF^P15A^. For DWORF^WT^, the *Z*-axis distributions of the side chains confirm the high dynamics of the unstructured N-terminal residues that transition between lipid-buried and solvent-exposed positions (**Figure 5A**). In both ensembles, Phe9 anchor the short amphipathic helix to the membrane, with its aromatic sidechain deeply buried in the acyl layer of the lipids. Polar side-chains of the N-terminal residues (Glu3, Lys4, Ser7, Thr8, Ser10, and His11) partition at the membrane/water interface, while Tyr31 and Ser35 side-chains stabilize the C-terminus inserting into the headgroup region of the luminal membrane leaflet. Specific lipid interactions (**Figure 5B**) involving Lys4 and His11 were persistent due to electrostatic interactions and hydrogen bonding with lipid phosphate groups. Based on topology-guided sequence alignments (**Figure 5C and 5D**), both Lys4 and His11 are predicted to bind SERCA, suggesting that these interactions with membranes could modulate DWORF binding to the ATPase (Winther et al., 2013, Akin et al., 2013). The interaction pattern between phospholipid headgroups and DWORF^P15A^ is similar to the wild-type, with N-terminal residues (1-7) showing a slight increase in lipid/side-chain contacts, and residues 10-15 showing only a slight decrease in the number and persistence of interactions (**Figure 5B**). The extensive interactions between the N-terminus cause heterogeneity in the upper leaflet as indicated by the larger variance in the phosphate *Z*-axis positions (1.0 Å) compared to the lower leaflet (0.38 Å) (**Figure 5E and 5F**). DWORF^WT^ causes localized defects in the upper membrane leaflet, which are due to domain Ib and domain Ia residues interacting with the surrounding lipids, while buried hydrophilic side-chains of Ser10 and His11 causes a localized thinning of the membrane (**Figure 5F**). DWORF^P15A^ causes more extensive deformations of the membrane, reducing its thickness by attracting the headgroups toward the hydrocarbon core of the lipids (**Figure 5F**). Lipid interactions at the C-terminus are driven by Tyr31 and Ser35. The aromatic side-chain of Tyr31 persistently snorkels to the lower leaflet to maximize solvent exposure of the hydroxyl group and hydrogen bonding with lipid phosphates.

**Figure 5:**
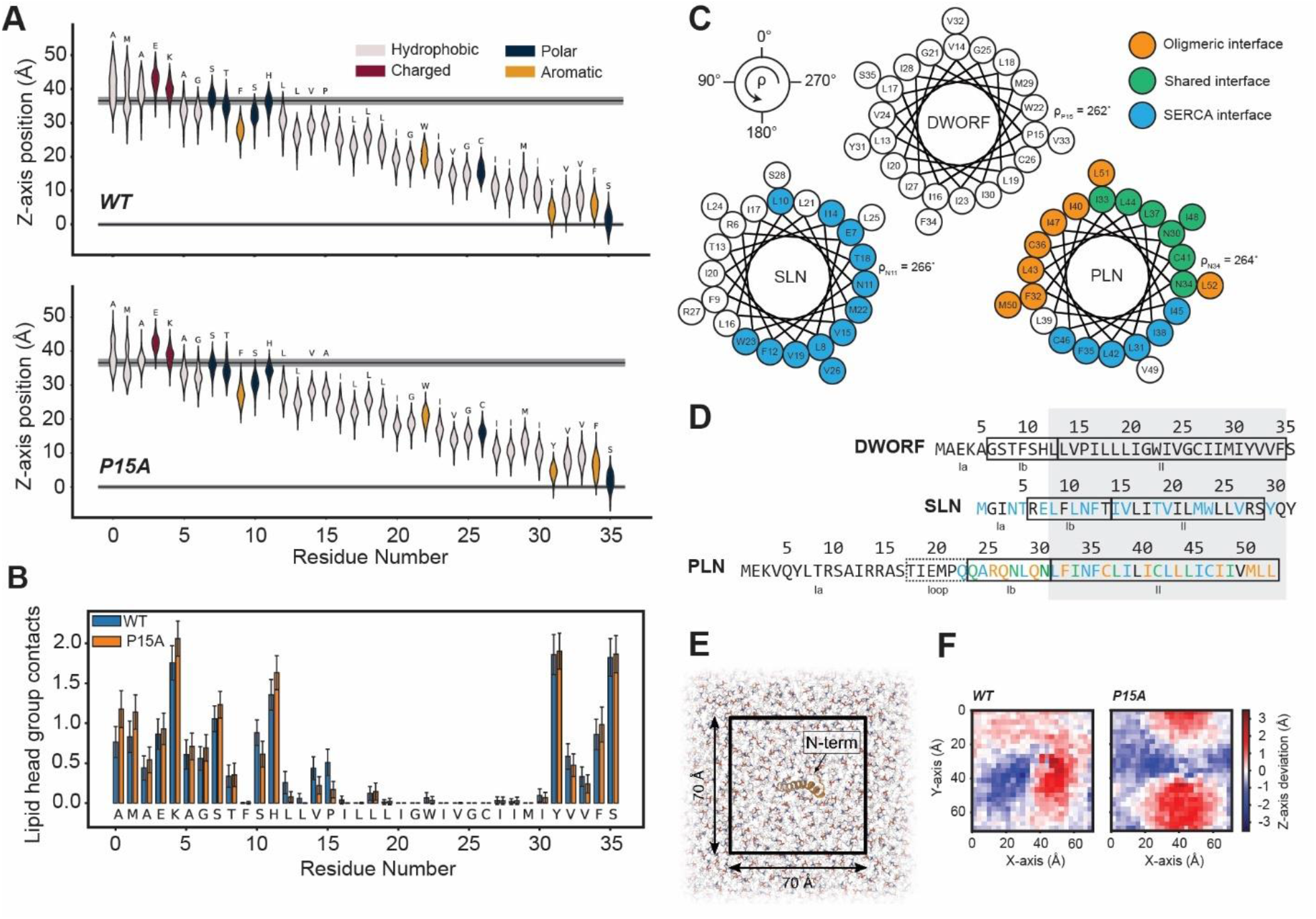
**(A)** Depth of insertion of DWORF (top) and DWORF^P15A^ (bottom) side-chains into the membrane. The vertical axis corresponds to the position of the center of mass of each side-chain in the *Z*-axis. Distributions of *Z*-axis positions across the ensemble are shown as violin shapes. The average positions of lipid phosphate atoms in each of the two leaflets are depicted as black horizontal lines (gray shades are the standard deviations). (**B)** Number of contacts established on average by each residue side-chain with nearby phospholipid head groups. **(C)** Helical wheel diagrams of TM domains of DWORF (this work), SLN (Mote et al., 2013, Wang et al., 2019) and PLN (Traaseth et al., 2009) rotated according to their azimuthal angles (ρ) reported from previously published SLF spectra (Mote et al., 2013, Wang et al., 2019, Traaseth et al., 2009). The oligomeric interface was determined by contact analysis of the PLN pentamer (Verardi et al., 2011). The SERCA binding interface was determined by intermolecular contacts in X-ray structures of the SERCA-SLN and SERCA-PLN complexes (Winther et al., 2013, Akin et al., 2013). **(D)** Sequence alignments based on topology. Residues of DWORF, SLN and PLN are colored according to their binding interface as per panel (C). **(E)** Local perturbation of the upper membrane leaflet by DWORF. A grid of 70 x 70 Å^2^ was extracted for analysis, where DWORF is positioned in the center and traverses the membrane from right to left along the X-axis (left panel). **(F)** The mean deviation of phosphate atoms from the average Z-position of the overall leaflet was calculated at each position of the grid, with negative and positive shifts representing the burial or uplifting of lipid headgroups, respectively.

During the RAOR-MD trajectory, the aromatic side chains of Phe9 and Phe34 are oriented with the plane of the aromatic ring perpendicular to bilayer surface, embedded into the acyl layer as anchor points for DWORF (**Figure 5A**). Interestingly, Trp22 located in the central region of domain II adopts an orientation horizontal to the membrane surface for approximately 90% of the conformations (**Figure 6**). This rotameric state of the Trp side chain is usually energetically unfavorable but preferred when positioned within the core of the membrane.

**Figure 6.**
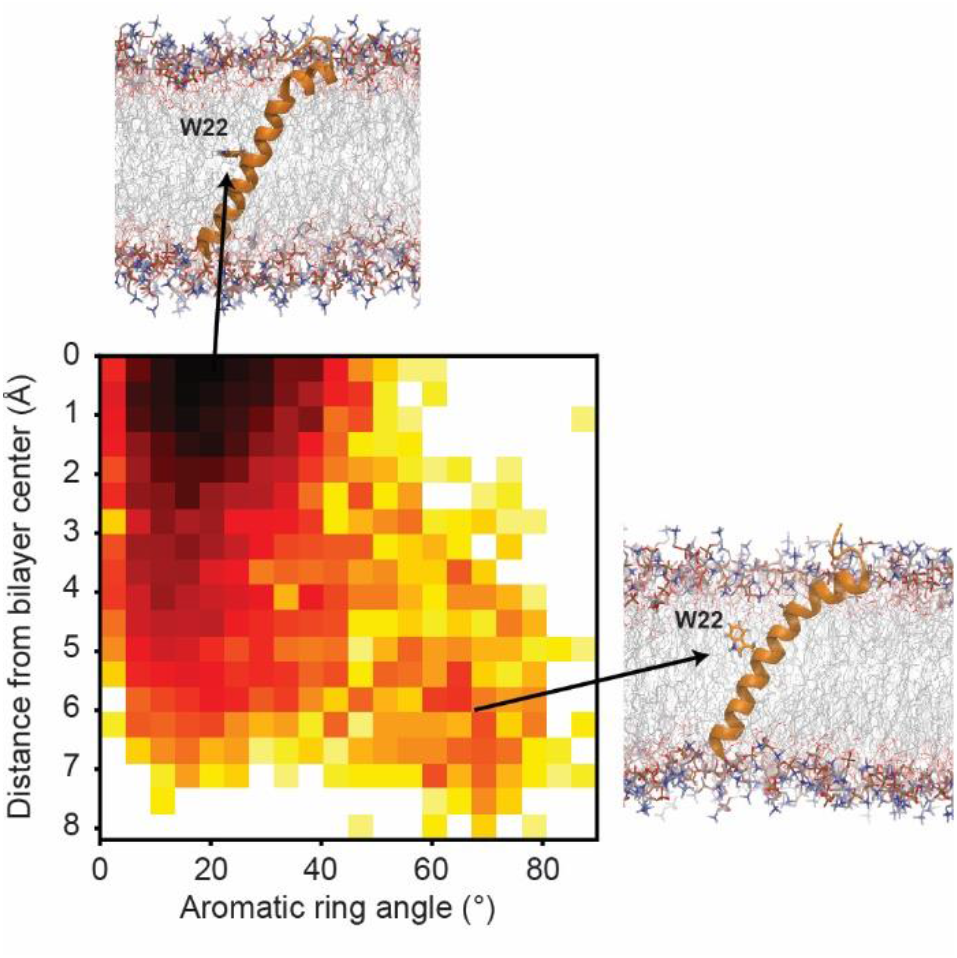
Distribution of Trp22 sidechain configurations. The horizontal axis shows the angle formed by the Trp22 aromatic ring normal vector and the membrane normal vector. The vertical shows the *Z*-component distance between the Trp22 side-chain centroid and the middle of the bilayer. A major and a minor configuration are depicted on the sides.

**Figure 7:**
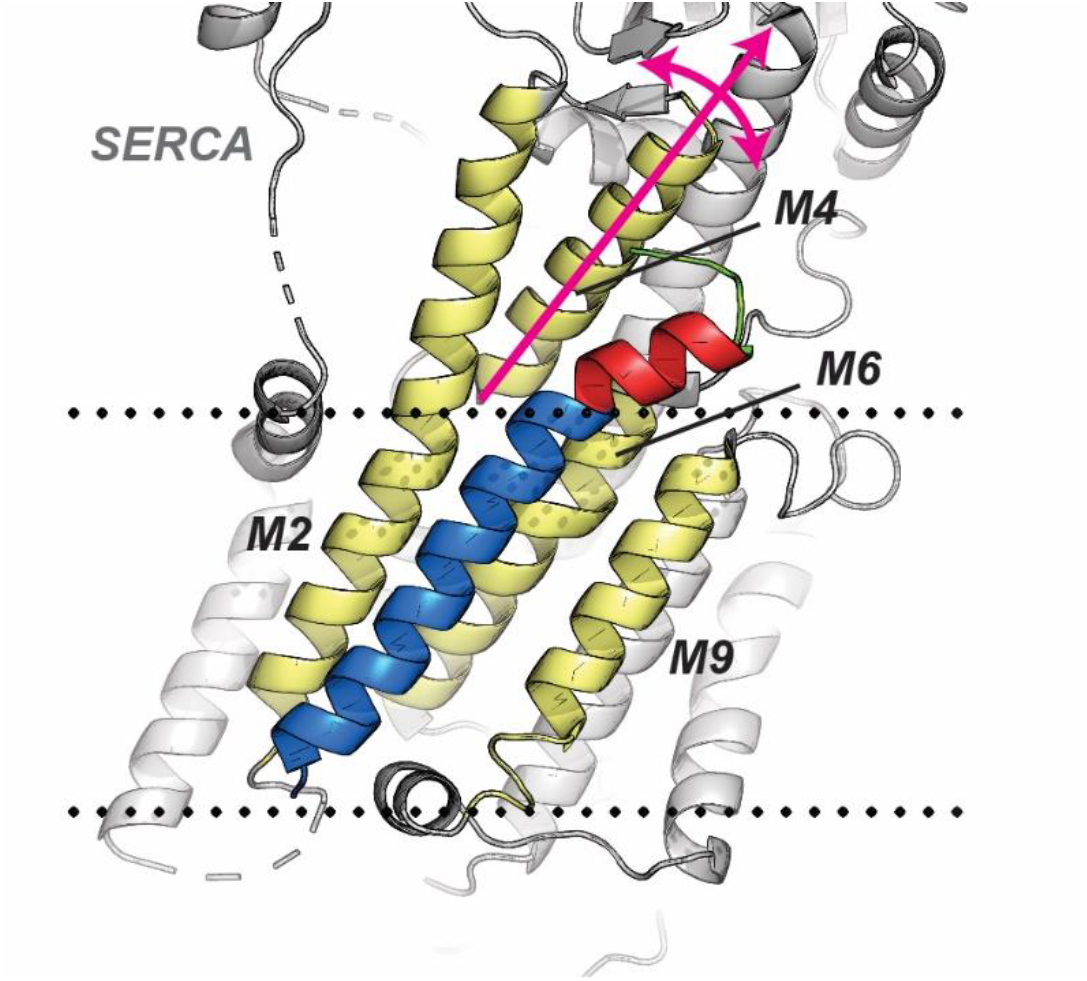
Model of DWORF bound to SERCA. Our lowest energy model of DWORF was manually docked in place of PLN in the X-ray structure of the SERCA-PLN complex (PDB 4Y3U). Docking was guided using the topologically based sequence alignment of Figure 5D. DWORF domains are colored green (Ia), red (Ib) and blue (II). Black dots indicate the membrane position determined using the PPM webserver (Lomize et al., 2012).

## DISCUSSION

SERCA’s regulatome has recently expanded from PLN and SLN to include DWORF, MLN, ELN and ALN (Nelson et al., 2016, Anderson et al., 2015, Payre and Desplan, 2016). Despite these bitopic membrane proteins having a relatively simple architecture, their effects on SERCA’s activity are highly diverse for reasons that are difficult to pinpoint by conventional structural methods. In this work, OS-ssNMR provided residue-specific structural and topological information for DWORF reconstituted into lipid bilayers. Like PLN and SLN, DWORF was subdivided into three distinct domains: a dynamic domain Ia, an amphipathic juxtamembrane domain Ib, and a hydrophobic TM domain II. Pro15, equivalent to the position of conserved Asn11 and Asn34 residues in SLN and PLN, respectively, induced a pronounced kink that distinguishes DWORF’s topology from the other regulators. Kinks, and other deviations from ideal geometry, are generally functionally important features in TM helices as they facilitate movement and introduce structural diversity into otherwise homogenous domains. Based on analogy to PLN and SLN, and their X-ray structures in complex with SERCA (Winther et al., 2013, Akin et al., 2013, Toyoshima et al., 2013), we surmise that DWORF’s domain Ib interacts with SERCA’s M4 helix. Since M4 undergoes large movements throughout SERCA’s enzymatic cycle (Dyla et al., 2020, Toyoshima, 2008) relative to the M2-M6-M9 binding groove, the kink could provide DWORF with an adaptive topology to better accommodate the structural transitions of the complex or even promote active states. Importantly, DWORF’s enhancement of SERCA enzymatic turnover rate (V_max_) is entirely abolished by a Pro15Ala mutation, but its inhibitory effect on Ca^2+^-binding affinity is retained, essentially becoming a typical inhibitor like the other regulators (Nelson et al., 2016). Interestingly, a kink, albeit far less pronounced, was still observed for the DWORF^P15A^ mutant, suggesting that the interactions between the cytoplasmic amphipathic helix and the lipid membrane contribute to the distortion of the helical structure (Yohannan et al., 2004).

Nelson et al. originally proposed a mechanism by which DWORF activates SERCA by displacing endogenous PLN from the binding groove and restoring Ca^2+^-binding affinity (pKCa) (Nelson et al., 2016). This indirect mechanism of activation was further supported by in-cell FRET studies (Makarewich et al., 2018). However, this was recently challenged by Li *et al*. who observed that DWORF directly activates SERCA’s pKCa without PLN having to be present (Li et al., 2021). In contrast to these studies, which used either animal-based or cell-based assays, our reconstituted systems found that DWORF directly activated SERCA’s V_max_ and did not outcompete PLN’s inhibitory effect on pK_Ca_. Rather, PLN and DWORF produced an overall weighted functional effect on SERCA. A similar increase in V_max_ was recently reported by Fisher *et al*., also using a reconstituted assay system, but an inhibitory effect on pK_Ca_ was not observed and competitive displacement of PLN was not determined (Fisher et al., 2020). These discrepancies are likely arising from variation in reconstitution protocols that affect PLN:SERCA and lipid:protein ratios.

Consistent with our work, recent FRET studies also found that the DWORF^P15A^ still bound SERCA, but a DWORF^P15A/W22A^ double mutation significantly reduced binding (Li et al., 2021). The central positioning of Trp22 in the membrane is rare since tryptophan is usually an interfacial anchoring residue to form hydrogen bonds and cation-π interaction with lipid headgroups (De Jesus and Allen, 2013). Indeed, Trp22 is predicted to lie in the middle of the SERCA-binding interface, based on its azimuthal angle, and RAOR-MD refinement suggested its side-chain occupied conformations that are atypical in lipid bilayers. RAOR-MD also modeled DWORF collective topological dynamics, specific lipid interactions and identified anchoring and snorkeling resides. In summary, OS-ssNMR and RAOR-MD refinement of DWORF reconstituted in lipid bilayers has provided critical high-resolution insights into the structural topology unpinning DWORF’s unique role as an endogenous activator SERCA.

## ACKNOWLEGMENTS

D.K.W. was supported by an American Heart Association Postdoctoral Fellowship (19POST34420009). This work was supported by the National Institutes of Health Heart Lung and Blood Institute grants HL143816 to S.L.R. and G.V. and HL092321 to S.L.R. We thank Caitlin Walker for her assistance with cloning the DWORF^P15A^ expression construct, and Dr. Maximo Sanz-Hernández for his contributions to the modelling work.

## STAR METHODS

### KEY RESOURCES TABLE

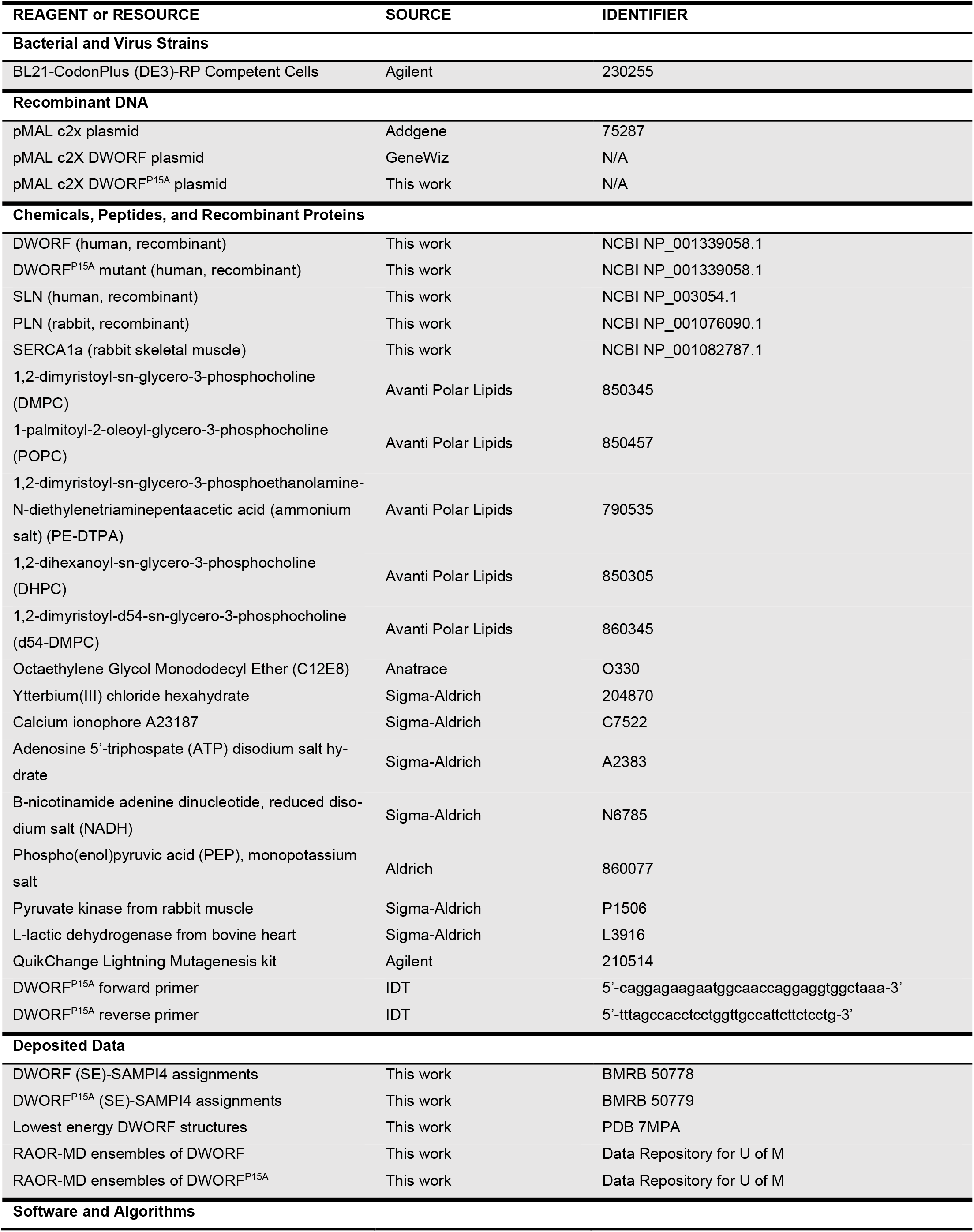
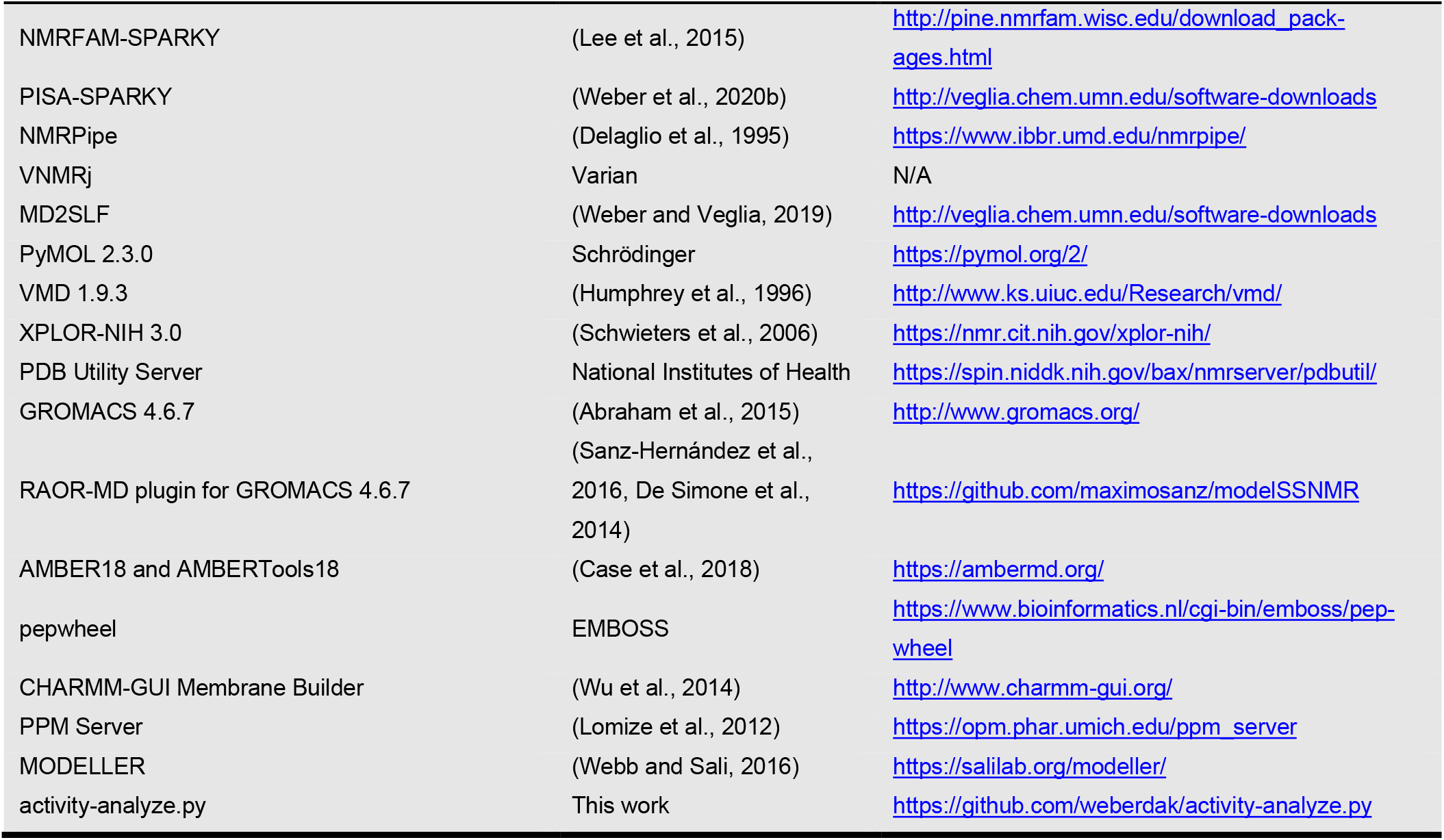

### RESOURCE AVAILABILITY

#### Lead Contact

Further information and requests for resources and reagents should be directed to and will be fulfilled by the lead contact, Gianluigi Veglia.

#### Materials Availability

Expression plasmids generated in this study are available on request.

#### Data and Code Availability

Assigned CSA-DC correlations from oriented (SE)-SAMPI4 spectra of DWORF and DWORF^P15A^, along raw and processed time-domain data, have been deposited on the Biological Magnetic Resonance Back with accession codes 50778 and 50779, respectively. Structures of DWORF obtained by XPOR-NIH simulated annealing calculations have been deposited in the RCSB Protein Data Bank (PDB 7MPA). RAOR-MD ensembles of DWORF and DWORF^P15A^ are available in the Data Repository for the University of Minnesota (DRUM; DOI pending).

### EXPERIMENTAL MODELS AND SUBJECT DETAILS

All experiments were caried out in vitro using DWORF obtained in this work either by recombinant expression or solid-phase peptide synthesis. Recombinant DWORF was expressed in *E. coli* BL21 (DE3) cells grown in minimal media required for uniform or selective ^15^N labelling. Synthetic DWORF for specific ^15^N-labelling was produced by solid-phase peptide synthesis.

## METHODS DETAILS

### Cloning, expression and purification of DWORF

The human DNA sequence for DWORF (NCBI Accession NP_001339058.1) was obtained from GeneWiz (South Plainfield, New Jersey) codon optimized for *E. coli* and ligated into a pMAL™ c2X expression vector (Ampicillin resistant; New England BioLabs) by EcoRI and HindIII restriction sites. The vector encoding DWORF fused to the N-terminus of the maltose binding protein (MBP) by a Tobacco Etch Virus (TEV) cleavage site (Glu-Asn-Leu-Tyr-Phe-Gln-Ala). Using this construct, the DWORF^P15A^ mutant was generated using a QuikChange Lightning Multi Site-Directed Mutagenesis Kit (Agilent) and primers 5’ −CAG GAG AAG AAT GGC AAC CAG GAG GTG GCT AAA −3’ and 5’-TTT AGC CAC CTC CTG GTT GCC ATT CTT CTC CTG −3’ (Integrated DNA Technologies).

Uniformly ^15^N-labelled DWORF was expressed as the soluble MBP-DWORF fusion protein using a similar approach as reported previously for PLN and SLN (Buck et al., 2003). A single colony of freshly transformed *E. coli* BL21-CodonPlus (DE3)-RP cells (Agilent) cells were used to inoculate overnight 1 L LB cultures (100 μg/mL ampicillin) at 28°C until an OD_600_ between 1.5 to 2.0, which were subsequently centrifuged (4,300 rpm, JS-4.3 rotor, 15 min) and resuspended into 1 L M9 minimal media (100 μg/mL ampicillin) to an OD_600_ of ∼0.75 with 1 g/L ^15^NH_4_Cl and 4 g/L D-glucose (D-glucose-^13^C_6_ for ^13^C,^15^N-labelled DWORF) as the sole nitrogen and carbon sources, respectively. Cultures were grown at 30°C to OD_600_ 1.0 then induced with 1 mM IPTG over 20 h to a final OD_600_ of ∼5. Cells were harvested by centrifuge (6,000 rpm, JLA-8.1000 rotor, 30 min) and stored at stored at −20°C (∼5.5 g wet cell mass per 1 L of M9). Selectively ^15^N-labelled DWORF was expressed from M9 media (free of NH_4_Cl) with 125 mg/L of the respective ^15^N-amino acid, 300 mg/L of non-scrambling and 450 mg/L scrambling-prone ^14^N-amino acids. Reverse-labeled DWORF was expressed in M9 minimal media (^15^NH_4_Cl) with 1 g/L of the respective ^14^N-labeled amino acid(s). Induction times for selective and reverse labeling growths were reduced to 3 to 4 h to limit scrambling.

For purification, cells were homogenized (Sorvall Omni Mixer) and lysed by sonification in 200 mL lysis buffer (20 mM sodium phosphate, 120 mM NaCl, 2 mM DTT, 1 mM EDTA, 0.1 mg/mL lysozyme, 0.5 % glycerol, 0.5% Tween 20 and a protease inhibitor cocktail, pH 7.3). The lysate was centrifuged (17,500 rpm, JA25.50 rotor, 4°C, 40 min) and the supernatant loaded onto a 30 mL bed volume of amylose resin (New England Biolabs). The resin was washed with 200 mL of buffer (20 mM sodium phosphate, 120 mM NaCl, pH 7.3) and eluted into 50 mL buffer including 50 mM maltose. Elution volumes were dialyzed overnight against 3 L of cleavage buffer (50 mM Tris-HCl, 2 mM β-mercaptoethanol, pH 7.3). All purification steps were done at 4°C and yielded up to 40 mg of fusion protein from 1 L M9 media.

MBP-DWORF was cleaved with recombinant TEV protease (3 mg per 100 mg MBP-DWORF) and 2 mM DTT for 3 h at 34°C to liberate insoluble DWORF (with an Ala0 residue), which was then pelleted by centrifugation (17,500 rpm, JA-25.50 rotor, 4°C, 20 min) and dissolved into 10% SDS and 50 mM DTT at approximately 4 mg/mL then stored at −20°C. DWORF was further purified by HPLC using a Vydac 214TP10154 250 x 4.6 mm 10-15 μm C4 column heated at 60°C and eluted using H_2_O/0.1% trifluoroacetic acid (TFA) and a linear gradient of isopropanol/0.1% TFA from 10 to 40% over 10 mins then to 80% over 50 min. The protein was lyophilized and confirmed by MALDI-MS (**Figure S1**). Typical yields were 4 mg of lyophilized DWORF from 1 L of growth.

### Solid-phase peptide synthesis of DWORF

Peptide synthesis was done on a CEM Liberty Blue solid phase peptide synthesizer at 0.05 mmole scale using Fmoc-Ser(tBu)-Wang resin LL (100-200 mesh, Novabiochem). After synthesis, the resin was collected and washed with DCM in a filtered cleavage syringe, then dried under vacuum to remove remaining solvent. DWORF was cleaved from the resin for 3 hrs in 3 to 5 mL of a solution composing of 20.6 mL of TFA, 1.25 g of phenol, 1.25 mL of thioanisol, 0.625 ml of ethanedithiol and 1.25 mL of ddH_2_O. The cleavage reaction was then solution filtered into ice cold diethyl ether and left one ice fore 10 min to precipitate the cleaved DWORF. DWORF was collected by centrifuge (4,000 rpm, JA-25.50 rotor, 4°C, 10 min). The pellet was washed twice more with ice cold diethyl ether and centrifugation, resuspended into water, the lyophilized. DWORF was then purified by HPLC as per the method used for recombinant protein.

### Reconstitution of DWORF into bicelles for OS-ssNMR spectroscopy

Long-chained lipids DMPC (29.0 mg) and POPC (8.2 mg) were cosolubilized from chloroform stocks then dried to a film with N_2_ followed by high-vacuum. Separately, PE-DTPA (1.0 mg) and DHPC (5.25 mg; *q* ratio of 4.4:1 to long-chained lipids) were cosolubilized and dried to a film with N_2_ and high vacuum. The DMPC and POPC film was resuspended into 150 μL of NMR buffer (20 mM HEPES, 20 mM KCl, 5 mM MgCl_2_, 2.5% glycerol, 0.02% NaN_3_, 5 mM EGTA, 5 mM DTT, pH 7.0) and freeze-thawed three times between liquid N_2_ and a 25°C water bath to obtain a cloudy liposome solution. The film of DHPC and PE-DTPA was resuspended separately into 120 uL of NMR buffer and used to solubilize lyophilized DWORF powder (1 to 3 mg) with vortex and pH-adjusted to 7.0 with KOH. The two solutions were then mixed at ice-cold temperature and vortexed while reaching room temperature. The process was repeated until long-chained lipids were completely solubilized. The mixture was a transparent liquid at ice-cold temperature and converted to a transparent gel at room temperature. The sample was placed back on ice and concentrated to ∼150 μL using a 0.5 mL 10 kDa MWCO centrifugal filter (Amicon) at 4°C. The sample was doped with 0.8 μL of 1 M YbCl_3_, corrected to pH 7.0 with KOH, then loaded into a 5 x 15 mm flat bottom sample cell (New Era NE-RG5-15-FB).

### Reconstitution of DWORF into liposomes for MAS-ssNMR spectroscopy

For MAS, 12 mg of d_54_-DMPC in chloroform was dried to a film using N_2_ gas and high vacuum, then resuspended into 500 µL of sample buffer (20 mM HEPES, 5 mM MgCl_2_, 20 mM KCl, 5 mM EGTA, 2.5% glycerol, 0.02% NaN_3_, 15 mM CaCl_2_, pH 7.2). 25 mg of C12E8 was added to form a clear solution of mixed micelles. DWORF, dissolved in 1 mL of 1% C12E8 detergent, was then combined with the mixed micelles and made to 2.5 mL with sample buffer. The solution was Incubated at 4°C and C12E8 removed by adding 1 g of BioBeads SM-2 (BioRad) in batches of 0.25 g added 30 minutes apart. After adding last batch, the further incubated 90 minutes at room temperature. The cloudy mixture of proteoliposomes was then removed from the BioBeads using a 25 G syringe and centrifuged (60,000 rpm, TLA-100.3 rotor, 4°C, 1 hr). The hydrated pellet was transferred directly to a 3.2 mm MAS rotor for NMR measurements.

### Oriented sample solid-state NMR spectroscopy

All oriented spectra were acquired on a Varian VNMRS spectrometer operating at a ^1^H frequency of 600 MHz and a BlackFox low-*E* static bicelle probe (Gor’kov et al., 2007). 1D ^15^N cross-polarization (CP)-based experiments used a 90° pulse length of 4 μs (62.5 kHz) on ^1^H and ^15^N channels with a contact time of 500 μs and 10% linear ramp on the ^1^H channel. Signals were detected with an acquisition time of 10 ms, 62.5 kHz SPINAL64 heteronuclear proton decoupling (Fung et al., 2000) and spectral width of 100 kHz in the direct dimension. The ^15^N transmitter frequency was set 162.5 ppm and externally referenced to ^15^NH_4_Cl at 39.3 ppm (Bertani et al., 2014). A recycle delay of 3 s was used.

2D separated local field (SLF) spectra were collected using a signal-enhanced (SE)-SAMPI4 experiment (Gopinath and Veglia, 2009, Gopinath et al., 2010a, Nevzorov and Opella, 2007). The indirect dipolar dimension utilized 25 increments and a scaled spectral width of 15.625 kHz. The *t*_1_ evolution period utilized ^1^H homonuclear decoupling with an RF field of 62.5 kHz and 96 μs dwell time, and phase-switched spin-lock pulses on ^15^N of 62.5 kHz. The sensitivity enhancement block used a *τ* delay of 75 μs and three cycles of phase-modulated Lee-Goldberg (PMLG) homonuclear decoupling (Vinogradov et al., 1999) with an effective RF field of 80 kHz. (SE)-SAMPI4 spectra were processed using NMRPipe (Delaglio et al., 1995) with a Lorentz-to-Gauss window function (150 Hz inverse exponential and 180 Hz Gaussian broadening widths) applied to ^15^N dimension and 2048 points of zero filling. The ^1^H-^15^N dipolar dimension was processed with a 90°-shifted sine bell window function and 1 kHz circular shift. Spectra were analyzed using NMRFAM-SPARKY (Lee et al., 2014) with PISA wheels fit using PISA-SPARKY plugin (Weber et al., 2020b). Tilt angles were fit by an exhaustive search of tilt and azimuthal angles in increments of 0.1 and 1°, respectively. The order parameter was fixed to 0.8 to reduce search space and prevent relatively disordered residues from systematically biasing fits. Errors quoted in the main text are the standard deviations of 20 replicate fits with peak positions randomly adjust within linewidths of 3.4 ppm and 0.55 kHz in the ^15^N CSA ^1^H-^15^N DC dimensions, respectively.

### Magic-angle-spinning solid-state NMR spectroscopy

MAS-ssNMR experiments were acquired using a Varian VNMRS spectrometer operating at a ^1^H frequency of 600 MHz and equipped with a Varian BioMAS 3.2 mm probe. A 90° pulse length of 2.5 μs was applied to the ^1^H channel and 6 μs to ^13^C and ^15^N channels. DARR and NCA spectra were acquired using DUMAS (Gopinath and Veglia, 2012) and TEDOR-NCA (Gopinath et al., 2019) polarization-optimized multi-acquisition experiments, respectively, applying 12.5 kHz spinning at 2 °C and recycle delays of 2 s. For SIM-CP matching conditions, an RF field of 35 kHz was applied on ^13^C and ^15^N, and a 60 kHz linearly ramped (90-100%) on ^1^H. The DARR experiment (Takegoshi et al., 2001) within the DUMAS sequence utilized a mixing time of 100 ms, 175 increments in the t_1_ dimension with 33.333 kHz spectral width and 320 scans. For the TEDOR-NCA experiment, a TEDOR (Jaroniec et al., 2002) mixing time of 1.28 ms was applied along with 3 ms mixing time for SPECIFIC-CP (Baldus et al., 1998) with RF fields of 19 kHz on ^13^C, 33 kHz on ^15^N and 100 kHz on ^1^H, and 2048 scans. For both the TEDOR and NCA elements, 40 t_1_ increments were sampled over a spectral width of 3.125 kHz. For both the DUMAS and TEDOR-NCA experiments, SPINAL ^1^H decoupling (Fung et al., 2000) was during t_1_ and t_2_, with a t_2_ acquisition time of 15 ms. For TEDOR-NCX spectra, a ^15^N t_1_ spectral width of 3.125 kHz was used. DARR spectra were processed using NMRPipe (Delaglio et al., 1995) with a Lorentz-to-Gauss window function (80 Hz inverse exponential and 120 Hz Gaussian broadening widths) applied to both ^13^C dimension and 32,000 points of zero filling. TEDOR spectra were processed using NMRPipe (Delaglio et al., 1995) with a with exponential multiplication of 150 Hz for both ^13^C and ^15^N dimension and 32,000 points of zero filling.

The ^13^C refocused INEPT TOBSY (RI-TOBSY) (Gopinath and Veglia, 2018) was acquired at 25 °C under 10 kHz MAS. An INEPT transfer period of 3.3 ms, ^13^C-^13^C TOBSY mixing time of 6.66 ms with P9_16_ recoupling (Hardy et al., 2001), and a ^13^C RF field of 30 kHz were used. WALTZ16 ^1^H decoupling (Shaka et al., 1983) was applied at 10 kHz during t_1_ and t_2_ evolution periods, with a t_2_ acquisition time of 20 ms at 100 KHz spectral width, and 100 t_1_ increments over a spectral width of 20 kHz. The TOBSY spectra were accumulated over 576 scans. TOSBY spectra were processed using NMRPipe (Delaglio et al., 1995) with exponential multiplication of 90 Hz for both ^13^C dimensions and 32,000 points of zero filling.

### Prediction of SLF spectra by unrestrained MD

The starting model of DWORF, truncated to residues Ser7 and Ser35 (DWORF_7-35_), was built as a continuous α-helix using PyMOL and oriented in an implicit membrane using the PPM Webserver (Lomize et al., 2012). DWORF_7-35_ was then embedded into a bilayer of 96 DMPC and 32 POPC lipids, with 150 mM KCl and 15 Å layer of water, using the CHARMM-GUI Membrane Builder (Jo et al., 2008, Wu et al., 2014). Simulations were run using CHARMM36 parameters and the pmemd.cuda module of the AMBER18 simulation package (Case et al., 2018). The simulation was run at 310 °K using the standard configurations supplied by CHARMM-GUI (as of July 2019). SLF spectra were predicted from the final 500 ns of a 600 ns unrestrained simulation using the MD2SLF Tcl script (http://veglia.chem.umn.edu/software-downloads) implemented in VMD 1.9.3 (Humphrey et al., 1996) as reported previously (Weber and Veglia, 2019). To reduce errors associated with slow-timescale tilt angle dynamics, DWORF was aligned throughout trajectory to a frame where it exhibited the approximate tilt angle estimated from an experimental (SE)-SAMPI4 spectrum (∼34°). An order parameter of 0.85 was applied to the results to approximate the rigid-body dynamics lost by aligning the trajectory. It is noteworthy that simulations were also done for the full length DWORF modelled entirely as an α-helix, but these did not converge to a properly formed SLF pattern and produced tilt angles much larger than experimentally indicated.

### Simulated annealing structure calculations

Starting structure of DWORF was built as an α-helix using the PDB Utility Server (Bax Group, National Institutes of Health). Simulated annealing calculations were run using XPLOR-NIH (Version 3.0) (Schwieters et al., 2006). An α-helical structure for residues Gly6 to Phe34 was enforced by restraining ϕ and ψ dihedral restraints within a range of 30° from ideal angles of −63° and −42°, respectively. The dihedrals angles connecting Leu12 and Leu13 (i.e., at the kink) were left unrestrained. The helical structure was also reinforced using 1-4 N-H-O hydrogen bond restraints hydrogen bond restraints and knowledge-based terms (Grishaev and Bax, 2004). Hydrogen bonding restraints were not applied between domain Ib and domain II residues.

_15_N CSA restraints (*δ*_*restraint*_) were taken directly from (SE)-SAMPI4 cross peaks (*δ*_*oriented*_) and modified to negate the isotropic chemical shift (*δ*_*iso*_) and scaled by the general order parameter (*S*) for protein and bicelle dynamics:

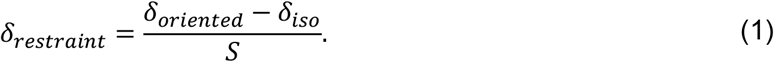

*δ*_*iso*_ was 122.2 ppm for non-glycine residues based on averaging the ^15^N shift tensor components of *δ*_11_ = 57.3 ppm, *δ*_22_ = 81.2 ppm and *δ*_33_ = 228.1 ppm (Murray et al., 2014), and 107.8 ppm for glycine based on components *δ*_11_ = 45.6 ppm, *δ*_22_ = 66.3 ppm and *δ*_33_ = 211.1 ppm (Straus et al., 2003). These tensors were implemented in XPLOR-NIH in the form *σ*_11_ = 64.9 ppm, *σ*_22_ = 41.0 ppm and *σ*_33_ = −105.9 ppm (non-glycine) and *σ*_11_ = 62.2 ppm, *σ*_22_ = 41.5 ppm and *σ*_33_ = −103.8 (glycine). An *S* of 0.8 was used as per PISA-wheel fitting. β angles associated with the non-glycine and glycine tensors were −17° and −21.6°, respectively. Similarly, ^1^H-^15^N DC restraints (*v*_*restraint*_) were scaled from SE-SAMPI4 cross peaks (*v*_*oriented*_) by:

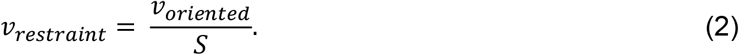

The maximum DC was fixed to 10.735 kHz (Denny et al., 2001) (*i*.*e*., *D*_*a*_ = 5.37). CSA and DC were applied as flat-well restraints with a total width of 10 ppm and 1 kHz, respectively.

The simulated annealing protocol included an initial 100 steps of Powell torsion angle minimization, followed by 20,000 steps of high temperature torsion dynamics at 3,500 K using the RepelPot repulsive non-bonded term (Nilges et al., 1988). Simulated annealing was then implemented from 3,500 K to 25 K in 12.5 K increments at 200 steps per increment. Dihedral force constants (CDIH) were ramped from 10 to 200 kcal mol^-1^ rad^-2^ during simulated annealing. The CSA force constant was ramped from 0.025 to 250 kcal mol^-1^ ppm^-2^ and DC constant from 0.075 to 750 kcal mol^-1^ kHz^-2^ (Shi et al., 2009). Statistical torsion-angle potentials (torsionDB) (Bermejo et al., 2012) were ramped from 0.02 to 2 kcal mol^-1^. The initial forces constants of the formally mentioned restraints were applied during high temperature dynamics stages. The Z-coordinates (*i*.*e*., depth-of-insertion) of the final structure was then optimized to an empirical virtual membrane model (Senes et al., 2007) with a hydrocarbon thickness of 25.7 Å to approximate the DMPC/POPC (4:1) bicelle composition (Marsh, 2013), followed by 500 steps of Powell torsion angle minimization then 500 steps of Powell cartesian minimization, applying restraints with full force constants. The 10 lowest energy structures (out of 5000 calculated) were then selected for further analysis and deposition into the Protein Data Bank. Structures having unrealistic relative orientations of domain Ia an Ib, and depth of insertions, which are an artifact of not having a membrane potential applied during the simulated annealing stage, were discarded from the final bundle of structures.

### Activity assays

SERCA1a was purified from the skeletal muscle of New Zealand white rabbits as previously described (Stokes and Green, 1990). For enzyme-couple activity assays (Reddy et al., 2003), SERCA was reconstituted, with and without a 2:1 molar ratio of regulators DWORF, DWORF^P15A^, PLN (wild-type rabbit), PLN^pS16^, or SLN (human), into DOPC/DOPE (4:1) bilayers at a lipid:SERCA molar ratio of 2500:1. These regular expressed using a similar protocol the DWORF, with PLN phosphorylated by protein-kinase A prior to cleavage of the MBP-PLN fusion protein. Reconstitution was achieved by co-solubilizing SERCA, regulators and lipids into C12E8 detergent (4:1 molar ratio to lipid), followed by removal of the detergent using BioBeads SM-2 (BioRad) to form proteoliposomes. Incubations with BioBeads were done at 4°C for 2 hrs followed by 1 hr at room temperature. Proteoliposomes were prepared to EGTA-buffered calcium concentrations ranging between 10^−8^ and 10^−5^ M (12 wells per assay, in triplicate) in a well composition of 40 nM SERCA, 100 μM lipids, 50 mM MOPS, 100 mM KCl, 5 mM MgCl_2_, 0.2 mM NADH, 0.5 mM PEP, 10 U/mL pyruvate kinase, 10 U/mL lactate dehydrogenase, and 7 μM calcium ionophore A23187 at pH 7.0. Reactions were initiated by adding 0.5 mM ATP and activity determined from the pseudo-first-order decay in NADH absorbance at 340 nm measured every 10 seconds for 10 minutes. Assays were done in triplicate at 25°C using a SpectroMax Plus 384 plate reader. Rates (activity) as a function of Ca^2+^ concentration (pCa) were fitted to a Hill function using the activity-analyze.py python script (https://github.com/weberdak/activity-analyze.py):

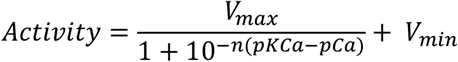

### Refinement by replica-averaged orientational-restrained molecular dynamics

The lowest-energy structure computed by XPLOR-NIH was initially equilibrated for 100 ns in an all-atom solvated membrane containing 75% DMPC, 25% POPC, using the CHARMM36m force field (Huang et al., 2017) and Tip3p waters (Jorgensen et al., 1983) with box of dimensions 7.0 × 7.0 × 7.3 nm^3^. An initial unrestrained sampling of a further 100 ns was run from which 8 structures were extracted to proceed with the replica-averaged refinement. Replica-averaged orientational-restrained molecular dynamics (RAOR-MD) were carried out as described previously (De Simone et al., 2014, Sanz-Hernández et al., 2016) using 8 parallel replicas. Restraining forces were introduced over 20 ns of equilibration, during which the force constants were linearly increased to reach 1250 J mol^-1^ ppm^-2^ and 18.5 J mol^-1^ kHz^-2^ for CSA and DC respectively. After the force equilibration, each replica was sampled for 80 ns. A final representative ensemble of 5,000 structures was extracted for analysis from the sampling phase.

To refine DWORF^P15A^, the mutation was introduced to the initial structure of DWORF using MOD-ELLER software (Webb and Sali, 2016). The resulting structure was embedded into the same membrane as WT DWORF and solvated in a box with the same dimensions. The DWORF^P15A^ structure was equilibrated without restraints for 200 ns, during which the initial kink in the structure was reduced due to the disappearance of the proline residue. Similar to WT DWORF, 8 structures were extracted from the last 100 ns of sampling in order to proceed with the RAOR-MD. RAOR-MD sampling was carried out using identical parameters as for WT.

All RAOR-MD samplings were performed with a timestep of 2 fs. Temperature was coupled using the V-rescale algorithm at 300 K (Bussi et al., 2007) and pressure using the Berendsen algorithm at 1 bar (Berendsen et al., 1984). The LINCS algorithm (Hess et al., 1997) was used for constrained and electrostatics interactions were calculated using the Particle-Mesh Ewald method (Darden et al., 1993).

## References

Abraham, M. J., Murtola, T., Schulz, R., PáLl, S., Smith, J. C., Hess, B. & Lindahl, E. 2015. GROMACS: High performance molecular simulations through multi-level parallelism from laptops to supercomputers. SoftwareX, 1-2, 19–25.

Akin, B. L., Hurley, T. D., Chen, Z. & Jones, L. R. 2013. The Structural Basis for Phospholamban Inhibition of the Calcium Pump in Sarcoplasmic Reticulum. Journal of Biological Chemistry, 288, 30181–30191.

Anderson, D. M., Anderson, K. M., Chang, C. L., Makarewich, C. A., Nelson, B. R., Mcanally, J. R., Kasaragod, P., Shelton, J. M., Liou, J., Bassel-Duby, R. & Olson, E. N. 2015. A micropeptide encoded by a putative long noncoding RNA regulates muscle performance. Cell, 160, 595–606.

Anderson, D. M., Makarewich, C. A., Anderson, K. M., Shelton, J. M., Bezprozvannaya, S., Bassel-Duby, R. & Olson, E. N. 2016. Widespread control of calcium signaling by a family of SERCA-inhibiting micropeptides. Science Signaling, 9, ra119–ra119.

Baldus, M., Petkova, A. T., Herzfeld, J. & Griffin, R. G. 1998. Cross polarization in the tilted frame: Assignment and spectral simplification in heteronuclear spin systems. Molecular Physics, 95, 1197–1207.

Berendsen, H. J. C., Postma, J. P. M., Van Gunsteren, W. F., Dinola, A. & Haak, J. R. 1984. Molecular dynamics with coupling to an external bath. Journal of Chemical Physics, 81, 3684–3690.

Bermejo, G. A., Clore, G. M. & Schwieters, C. D. 2012. Smooth statistical torsion angle potential derived from a large conformational database via adaptive kernel density estimation improves the quality of NMR protein structures. Protein Science, 21, 1824–1836.

Bers, D. M. 2002. Cardiac excitation–contraction coupling. Nature, 415, 198–205.

Bertani, P., Raya, J. & Bechinger, B. 2014. 15N chemical shift referencing in solid state NMR. Solid State Nuclear Magnetic Resonance, 61–62, 15-18.

Buck, B., Zamoon, J., Kirby, T. L., Desilva, T. M., Karim, C., Thomas, D. & Veglia, G. 2003. Overexpression, purification, and characterization of recombinant Ca-ATPase regulators for high-resolution solution and solid-state NMR studies. Protein Expression and Purification, 30, 253–261.

Bussi, G., Donadio, D. & Parrinello, M. 2007. Canonical sampling through velocity rescaling. Journal of Chemical Physics, 126, 014101–014101.

Case, D. A., Ben-Shalom, I. Y., Brozell, S. R., Cerutti, D. S., Cheatham Iii, T. E., Cruzeiro, V. W. D., Darden, T. A., Duke, R. E., Ghoreishi, D., Gilson, M. K., Gohlke, H., Goetz, A. W., Greene, D., Harris, R., Homeyer, N., Izadi, S., Kovalenko, A., Kurtzman, T., Lee, T. S., Legrand, S., Li, P., Lin, C., Liu, J., Luchko, T., Luo, R., Mermelstein, D. J., Merz, K. M., Miao, Y., Monard, G., Nguyen, C., Nguyen, H., Omelyan, I., Onufriev, A., Pan, F., Qi, R., Roe, D. R., Roitberg, A., Sagui, C., Schott-Verdugo, S., Shen, J., Simmerling, C. L., Smith, J., Salomon-Ferrer, R., Swails, J., Walker, R. C., Wang, J., Wei, H., Wolf, R. M., Wu, X., Xiao, L., York, D. M. & Kollman, P. A. 2018. AMBER18. San Francisco: University of California.

Darden, T., York, D. & Pedersen, L. 1993. Particle mesh Ewald: An N°log(N) method for Ewald sums in large systems. Journal of Chemical Physics, 98, 10089–10092.

De Jesus, A. J. & Allen, T. W. 2013. The role of tryptophan side chains in membrane protein anchoring and hydrophobic mismatch. Biochimica et Biophysica Acta - Biomembranes, 1828, 864–876.

De Simone, A., Mote, K. R. & Veglia, G. 2014. Structural Dynamics and Conformational Equilibria of SERCA Regulatory Proteins in Membranes by Solid-State NMR Restrained Simulations. Biophysical Journal, 106, 2566–2576.

Delaglio, F., Grzesiek, S., Vuister, G. W., Zhu, G., Pfeifer, J. & Bax, A. 1995. NMRPipe: A multidimensional spectral processing system based on UNIX pipes. Journal of Biomolecular NMR, 6, 277–293.

Denny, J. K., Wang, J., Cross, T. A. & Quine, J. R. 2001. PISEMA powder patterns and PISA wheels. Journal of Magnetic Resonance, 152, 217–226.

Dyla, M., KjÆRgaard, M., Poulsen, H. & Nissen, P. 2020. Structure and Mechanism of P-Type ATPase Ion Pumps. Annual Review of Biochemistry, 89, 583–603.

Fisher, M. E., Bovo, E., Cho, E. E., Pribadi, M. P., Dalton, M. P., Lemieux, M. J., Rathod, N., Aguayo-Ortiz, R., Espinoza-Fonseca, L. M., Robia, S. L., Zima, A. V. & Young, H. S. 2020. Dwarf open reading frame (DWORF) peptide is a direct activator of the sarcoplasmic reticulum calcium pump SERCA. bioRxiv, 53, 1689–1699.

Fung, B. M., Khitrin, A. K. & Ermolaev, K. 2000. An Improved Broadband Decoupling Sequence for Liquid Crystals and Solids. Journal of Magnetic Resonance, 142, 97–101.

Gamu, D., Juracic, E. S., Hall, K. J. & Tupling, A. R. 2020. The sarcoplasmic reticulum and SERCA: a nexus for muscular adaptive thermogenesis. Applied Physiology, Nutrition, and Metabolism, 45, 1–10.

Gopinath, T. & Veglia, G. 2009. Sensitivity Enhancement in Static Solid-State NMR Experiments via Single-and Multiple-Quantum Dipolar Coherences. Journal of the American Chemical Society, 131, 5754–5756.

Gopinath, T. & Veglia, G. 2012. Dual acquisition magic-angle spinning solid-State NMR-spectroscopy: Simultaneous acquisition of multidimensional spectra of biomacromolecules. Angewandte Chemie - International Edition, 51, 2731–2735.

Gopinath, T. & Veglia, G. 2018. Probing membrane protein ground and conformationally excited states using dipolar-and J-coupling mediated MAS solid state NMR experiments. Methods, 148, 115–122.

Gopinath, T., Verardi, R., Traaseth, N. J. & Veglia, G. 2010a. Sensitivity Enhancement of Separated Local Field Experiments: Application to Membrane Proteins. The Journal of Physical Chemistry B, 114, 5089–5095.

Gopinath, T., Verardi, R., Traaseth, N. J. & Veglia, G. 2010b. Sensitivity enhancement of separated local field experiments: Application to membrane proteins. Journal of Physical Chemistry B, 114, 5089–5095.

Gopinath, T., Wang, S., Lee, J., Aihara, H. & Veglia, G. 2019. Hybridization of TEDOR and NCX MAS solid-state NMR experiments for simultaneous acquisition of heteronuclear correlation spectra and distance measurements. Journal of Biomolecular NMR, 73, 141–153.

Gor’Kov, P. L., Chekmenev, E. Y., Li, C., Cotten, M., Buffy, J. J., Traaseth, N. J., Veglia, G. & Brey, W. W. 2007. Using low-E resonators to reduce RF heating in biological samples for static solid-state NMR up to 900 MHz. Journal of Magnetic Resonance, 185, 77–93.

Grishaev, A. & Bax, A. 2004. An Empirical Backbone−Backbone Hydrogen-Bonding Potential in Proteins and Its Applications to NMR Structure Refinement and Validation. Journal of the American Chemical Society, 126, 7281–7292.

Hardy, E. H., Verel, R. & Meier, B. H. 2001. Fast MAS total through-bond correlation spectroscopy. Journal of Magnetic Resonance, 148, 459–464.

Hess, B., Bekker, H., Berendsen, H. J. C. & Fraaije, J. G. E. M. 1997. LINCS: A linear constraint solver for molecular simulations. Journal of Computational Chemistry, 18, 1463–1472.

Huang, J., Rauscher, S., Nawrocki, G., Ran, T., Feig, M., De Groot, B. L., GrubmÜLler, H. & Mackerell, A. D. 2017. CHARMM36m: an improved force field for folded and intrinsically disordered proteins. Nature Methods, 14, 71–73.

Humphrey, W., Dalke, A. & Schulten, K. 1996. VMD: Visual molecular dynamics. Journal of Molecular Graphics, 14, 33–38.

Jaroniec, C. P., Filip, C. & Griffin, R. G. 2002. 3D TEDOR NMR experiments for the simultaneous measurement of multiple carbon-nitrogen distances in uniformly 13C,15N-labeled solids. Journal of the American Chemical Society, 124, 10728–10742.

Jo, S., Kim, T., Iyer, V. G. & Im, W. 2008. CHARMM-GUI: A web-based graphical user interface for CHARMM. Journal of Computational Chemistry, 29, 1859–1865.

Jorgensen, W. L., Chandrasekhar, J., Madura, J. D., Impey, R. W. & Klein, M. L. 1983. Comparison of simple potential functions for simulating liquid water. Journal of Chemical Physics, 79, 926–935.

Kim, S., Quine, J. R. & Cross, T. A. 2001. Complete cross-validation and R-factor calculation of a solid-state NMR derived structure. Journal of the American Chemical Society, 123, 7292–7298.

Kranias, E. G. & Bers, D. M. 2007. Calcium and cardiomyopathies, Dordrecht, Springer Netherlands.

Kranias, E. G. & Hajjar, R. J. 2017. The Phospholamban Journey 4 Decades After Setting Out for Ithaka. Circulation Research, 120, 781–783.

Lee, W., Tonelli, M. & Markley, J. L. 2014. NMRFAM-SPARKY: enhanced software for biomolecular NMR spectroscopy. Bioinformatics, 31, 1325–1327.

Lee, W., Tonelli, M. & Markley, J. L. 2015. NMRFAM-SPARKY: enhanced software for biomolecular NMR spectroscopy. Bioinformatics, 31, 1325–1327.

Li, A., Yuen, S. L., Stroik, D. R., Kleinboehl, E., Cornea, R. L. & Thomas, D. D. 2021. The transmembrane peptide DWORF activates SERCA2a via dual mechanisms. Journal of Biological Chemistry.

Lomize, M. A., Pogozheva, I. D., Joo, H., Mosberg, H. I. & Lomize, A. L. 2012. OPM database and PPM web server: Resources for positioning of proteins in membranes. Nucleic Acids Research, 40, D370–D376.

Luo, M. & Anderson, M. E. 2013. Mechanisms of Altered Ca<sup>2+</sup> Handling in Heart Failure. Circulation Research, 113, 690–708.

Maclennan, D. H. & Kranias, E. G. 2003. Phospholamban: a crucial regulator of cardiac contractility. Nature reviews: Molecular cell biology, 4, 566–577.

Makarewich, C. A. 2020. The hidden world of membrane microproteins. Experimental Cell Research, 388, 111853–111853.

Makarewich, C. A., Munir, A. Z., Schiattarella, G. G., Bezprozvannaya, S., Raguimova, O. N., Cho, E. E., Vidal, A. H., Robia, S. L., Bassel-Duby, R. & Olson, E. N. 2018. The DWORF micropeptide enhances contractility and prevents heart failure in a mouse model of dilated cardiomyopathy. eLife, 7, 1–23.

Marassi, F. M. & Opella, S. J. 2000. A Solid-State NMR Index of Helical Membrane Protein Structure and Topology. Journal of Magnetic Resonance, 144, 150–155.

Marsh, D. 2013. Handbook of Lipid Bilayers, CRC Press.

Morales Rodriguez, B., DomíNguez-RodríGuez, A., Benitah, J.-P., Lefebvre, F., Marais, T., Mougenot, N., Beauverger, P., Bonne, G., Briand, V., GóMez, A.-M. & Muchir, A. 2020. Activation of sarcolipin expression and altered calcium cycling in LMNA cardiomyopathy. Biochemistry and Biophysics Reports, 22, 100767–100767.

Mote, K. R., Gopinath, T. & Veglia, G. 2013. Determination of structural topology of a membrane protein in lipid bilayers using polarization optimized experiments (POE) for static and MAS solid state NMR spectroscopy. Journal of Biomolecular NMR, 57, 91–102.

Murray, D. T., Hung, I. & Cross, T. A. 2014. Assignment of oriented sample NMR resonances from a three transmembrane helix protein. Journal of Magnetic Resonance, 240, 34–44.

Nelson, B. R., Makarewich, C. A., Anderson, D. M., Winders, B. R., Troupes, C. D., Wu, F., Reese, A. L., Mcanally, J. R., Chen, X., Kavalali, E. T., Cannon, S. C., Houser, S. R., Bassel-Duby, R. & Olson, E. N. 2016. A peptide encoded by a transcript annotated as long noncoding RNA enhances SERCA activity in muscle. Science, 351, 271–275.

Nevzorov, A. A. & Opella, S. J. 2007. Selective averaging for high-resolution solid-state NMR spectroscopy of aligned samples. Journal of Magnetic Resonance, 185, 59–70.

Nilges, M., Clore, G. M. & Gronenborn, A. M. 1988. Determination of three-dimensional structures of proteins from interproton distance data by hybrid distance geometry-dynamical simulated annealing calculations. FEBS Letters, 229, 317–324.

Palmer, B. F. & Clegg, D. J. 2017. Non-shivering thermogenesis as a mechanism to facilitate sustainable weight loss. Obesity Reviews, 18, 819–831.

Payre, F. & Desplan, C. 2016. RNA. Small peptides control heart activity. Science (New York, N.Y.), 351, 226–7.

Periasamy, M., Maurya, S. K., Sahoo, S. K., Singh, S., Reis, F. C. G. & Bal, N. C. 2017. Role of SERCA pump in muscle thermogenesis and metabolism. Comprehensive Physiology, 7, 879–890.

Prosser, R. S., Volkov, V. B. & Shiyanovskaya, I. V. 1998. Novel Chelate-Induced Magnetic Alignment of Biological Membranes. Biophysical Journal, 75, 2163–2169.

Ramamoorthy, A., Wei, Y. & Lee, D.-K. 2004. PISEMA Solid-State NMR Spectroscopy. Annual Reports on NMR Spectroscopy. Academic Press.

Reddy, L. G., Cornea, R. L., Winters, D. L., Mckenna, E. & Thomas, D. D. 2003. Defining the Molecular Components of Calcium Transport Regulation in a Reconstituted Membrane System. Biochemistry, 42, 4585–4592.

Sanz-HernáNdez, M., Vostrikov, V. V., Veglia, G. & De Simone, A. 2016. Accurate Determination of Conformational Transitions in Oligomeric Membrane Proteins. Scientific Reports, 6, 23063–23063.

Schwieters, C. D., Kuszewski, J. J. & Marius Clore, G. 2006. Using Xplor-NIH for NMR molecular structure determination. Progress in Nuclear Magnetic Resonance Spectroscopy, 48, 47–62.

Senes, A., Chadi, D. C., Law, P. B., Walters, R. F. S., Nanda, V. & Degrado, W. F. 2007. Ez, a Depth-dependent Potential for Assessing the Energies of Insertion of Amino Acid Side-chains into Membranes: Derivation and Applications to Determining the Orientation of Transmembrane and Interfacial Helices. Journal of Molecular Biology, 366, 436–448.

Shaka, A. J., Keeler, J., Frenkiel, T. & Freeman, R. 1983. An improved sequence for broadband decoupling: WALTZ-16. Journal of Magnetic Resonance, 52, 335–338.

Shanmugam, M., Li, D., Gao, S., Fefelova, N., Shah, V., Voit, A., Pachon, R., Yehia, G., Xie, L.- H. & Babu, G. J. 2015. Cardiac Specific Expression of Threonine 5 to Alanine Mutant Sarcolipin Results in Structural Remodeling and Diastolic Dysfunction. PLOS ONE, 10, e0115822–e0115822.

Shi, L., Traaseth, N. J., Verardi, R., Cembran, A., Gao, J. & Veglia, G. 2009. A refinement protocol to determine structure, topology, and depth of insertion of membrane proteins using hybrid solution and solid-state NMR restraints. Journal of Biomolecular NMR, 44, 195–205.

Singh, D. R., Dalton, M. P., Cho, E. E., Pribadi, M. P., Zak, T. J., ŠEflová, J., Makarewich, C. A., Olson, E. N. & Robia, S. L. 2019. Newly Discovered Micropeptide Regulators of SERCA Form Oligomers but Bind to the Pump as Monomers. Journal of Molecular Biology, 431, 4429–4443.

Stokes, D. L. & Green, N. M. 1990. Three-dimensional crystals of CaATPase from sarcoplasmic reticulum. Symmetry and molecular packing. Biophysical Journal, 57, 1–14.

Straus, S. K., Scott, W. R. P. & Watts, A. 2003. Assessing the effects of time and spatial averaging in 15N chemical shift/15N-1H dipolar correlation solid state NMR experiments. Journal of biomolecular NMR, 26, 283–95.

Takegoshi, K., Nakamura, S. & Terao, T. 2001. 13C-1H dipolar-assisted rotational resonance in magic-angle spinning NMR. Chemical Physics Letters, 344, 631–637.

Toyoshima, C. 2008. Structural aspects of ion pumping by Ca2+-ATPase of sarcoplasmic reticulum. Archives of Biochemistry and Biophysics, 476, 3–11.

Toyoshima, C., Iwasawa, S., Ogawa, H., Hirata, A., Tsueda, J. & Inesi, G. 2013. Crystal structures of the calcium pump and sarcolipin in the Mg2+-bound E1 state. Nature, 495, 260–264.

Traaseth, N. J., Shi, L., Verardi, R., Mullen, D. G., Barany, G. & Veglia, G. 2009. Structure and topology of monomeric phospholamban in lipid membranes determined by a hybrid solution and solid-state NMR approach. Proceedings of the National Academy of Sciences, 106, 10165–10170.

Verardi, R., Shi, L., Traaseth, N. J., Walsh, N. & Veglia, G. 2011. Structural topology of phospholamban pentamer in lipid bilayers by a hybrid solution and solid-state NMR method. Proceedings of the National Academy of Sciences, 108, 9101–9106.

Vinogradov, E., Madhu, P. K. & Vega, S. 1999. High-resolution proton solid-state NMR spectroscopy by phase-modulated Lee-Goldburg experiment. Chemical Physics Letters, 314, 443–450.

Wang, J., Denny, J., Tian, C., Kim, S., Mo, Y., Kovacs, F., Song, Z., Nishimura, K., Gan, Z., Fu, R., Quine, J. R. & Cross, T. A. 2000. Imaging Membrane Protein Helical Wheels. Journal of Magnetic Resonance, 144, 162–167.

Wang, S., Gopinath, T. & Veglia, G. 2019. Improving the quality of oriented membrane protein spectra using heat-compensated separated local field experiments. Journal of Biomolecular NMR.

Webb, B. & Sali, A. 2016. Comparative Protein Structure Modeling Using MODELLER. Current Protocols in Protein Science, 86, 2.9.1-2.9.37.

Weber, D. K., Larsen, E. K., Gopinath, T. & Veglia, G. 2020a. Hybridizing isotropic and anisotropic solid-state NMR restraints for membrane protein structure determination. In: Separovic, F. & Sani, M.-A. (eds.). IOP Publishing.

Weber, D. K. & Veglia, G. 2019. A Theoretical Assessment of the Structure Determination of Multi-Span Membrane Proteins by Oriented Sample Solid-State NMR Spectroscopy. Australian Journal of Chemistry, 73, 246–251.

Weber, D. K., Wang, S., Markley, J. L., Veglia, G. & Lee, W. 2020b. PISA-SPARKY: an interactive SPARKY plugin to analyze oriented solid-state NMR spectra of helical membrane proteins. Bioinformatics, 1-2.

Williams, C. J., Headd, J. J., Moriarty, N. W., Prisant, M. G., Videau, L. L., Deis, L. N., Verma, V., Keedy, D. A., Hintze, B. J., Chen, V. B., Jain, S., Lewis, S. M., Arendall, W. B., Snoeyink, J., Adams, P. D., Lovell, S. C., Richardson, J. S. & Richardson, D. C. 2018. MolProbity: More and better reference data for improved all-atom structure validation. Protein Science, 27, 293–315.

Winther, A.-M. L., Bublitz, M., Karlsen, J. L., MØLler, J. V., Hansen, J. B., Nissen, P. & Buch-Pedersen, M. J. 2013. The sarcolipin-bound calcium pump stabilizes calcium sites exposed to the cytoplasm. Nature, 495, 265–9.

Wu, E. L., Cheng, X., Jo, S., Rui, H., Song, K. C., Davila-Contreras, E. M., Qi, Y., Lee, J., Monje-Galvan, V., Venable, R. M., Klauda, J. B. & Im, W. 2014. CHARMM-GUI Membrane Builder toward realistic biological membrane simulations. Journal of Computational Chemistry, 35, 1997–2004.

Yohannan, S., Faham, S., Yang, D., Whitelegge, J. P. & Bowie, J. U. 2004. The evolution of transmembrane helix kinks and the structural diversity of G protein-coupled receptors. Proceedings of the National Academy of Sciences, 101, 959–963.

Zheng, J., Yancey, D. M., Ahmed, M. I., Wei, C. C., Powell, P. C., Shanmugam, M., Gupta, H., Lloyd, S. G., Mcgiffin, D. C., Schiros, C. G., Denney, T. S., Babu, G. J. & Dell’Italia, L. J. 2014. Increased sarcolipin expression and adrenergic drive in humans with preserved left ventricular ejection fraction and chronic isolated mitral regurgitation. Circulation: Heart Failure, 7, 194–202.

Zhihao, L., Jingyu, N., Lan, L., Michael, S., Rui, G., Xiyun, B., Xiaozhi, L. & Guanwei, F. 2020. SERCA2a: a key protein in the Ca2+ cycle of the heart failure. Heart Failure Reviews, 25, 523–535.

