## Supplementary Figures and Table1 for "A Kink in DWORF Helical Structure Controls the Activation of the Sarco-plasmic Reticulum Ca^2+^-ATPase"

[*] To whom correspondence should be addressed:

### SUPPLEMENTAL INFORMATION

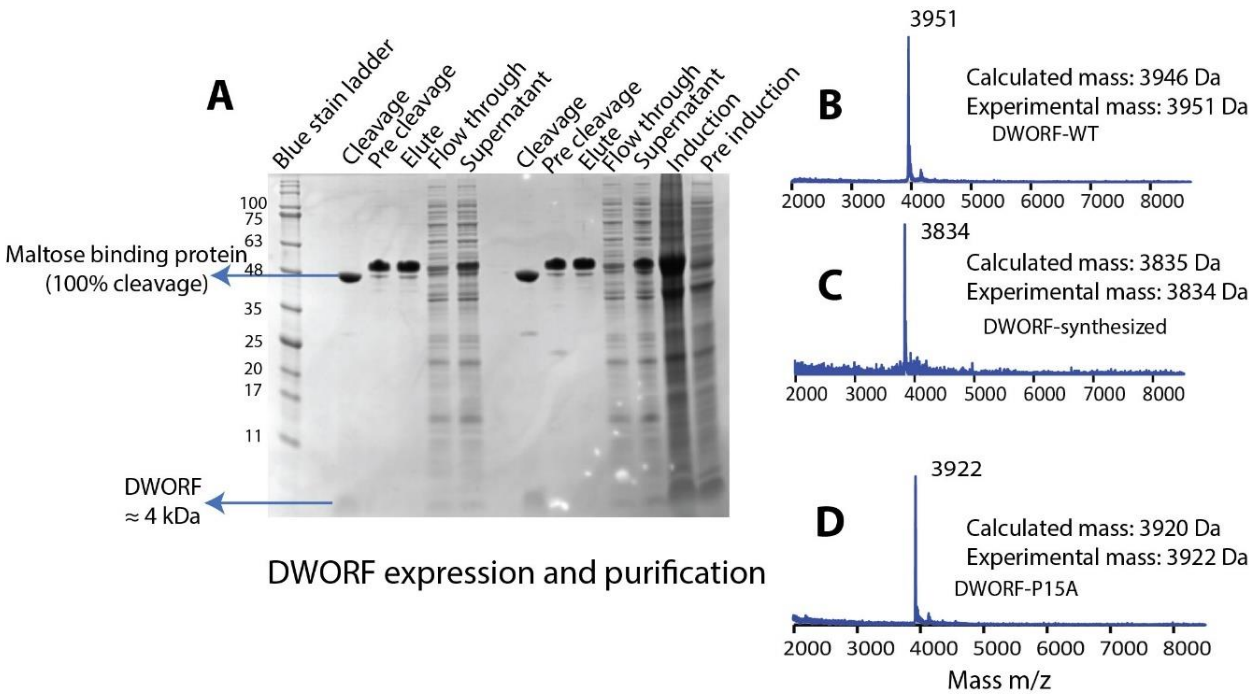


**Figure S1. (A)** SDS phase gel shows the expression and purification of DWORF. MALDI-MS of **(B)** $^{15}$N DWORF$^{WT}$, **(C)** DWORF synthesized by solid phase peptide synthesis, **(D)** and $^{15}$N DWORF$^{P15A}$.

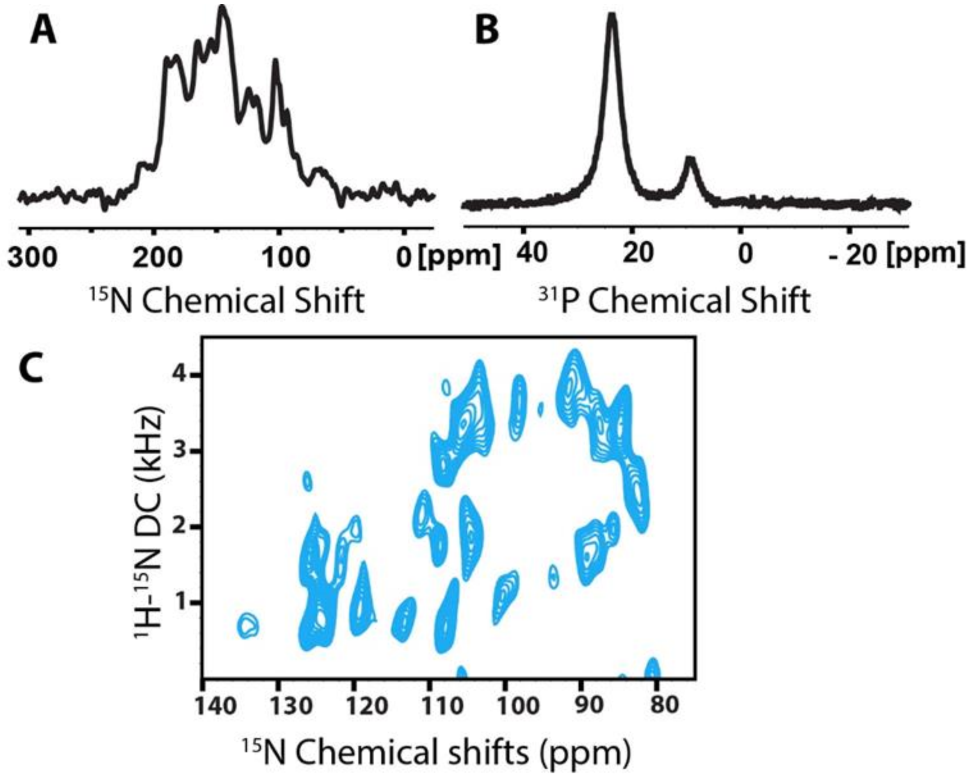


**Figure S2. (A)** $^{15}$N CP and **(B)** $^{31}$P NMR spectra of $^{15}$N DWORF reconstituted into *flipped* DMPC/POPC/PE-DTPA/DHPC bicelles. **(C)** (SE)-SAMPI4 spectrum of $^{15}$N DWORF in *unflipped* bicelles.

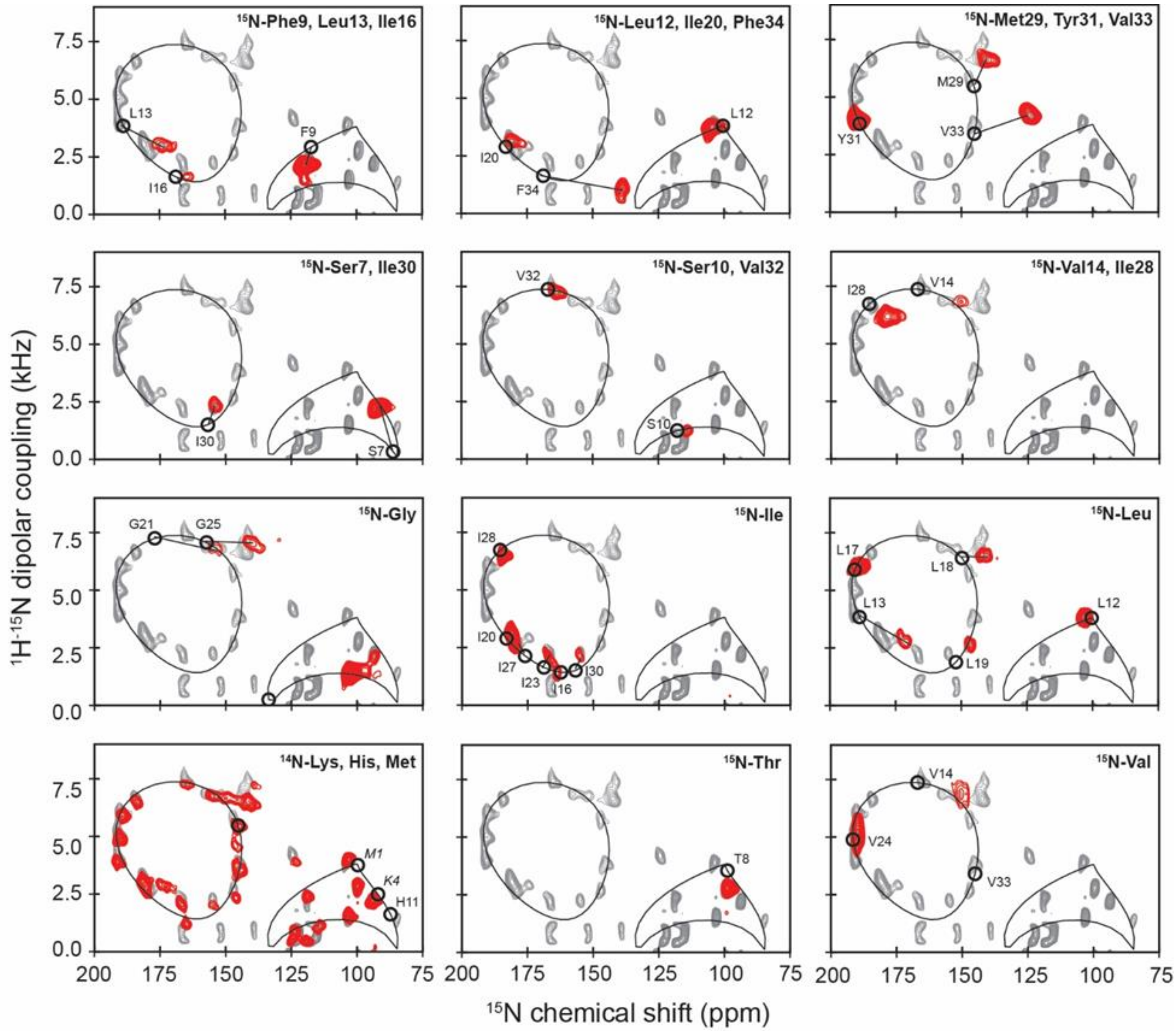


**Figure S3.** (SE)-SAMPI4 spectra of selectively labelled DWORF. Circles indicate theoretical peak positions from PISA-wheel fitting. Italicised labels specify undetected residues.

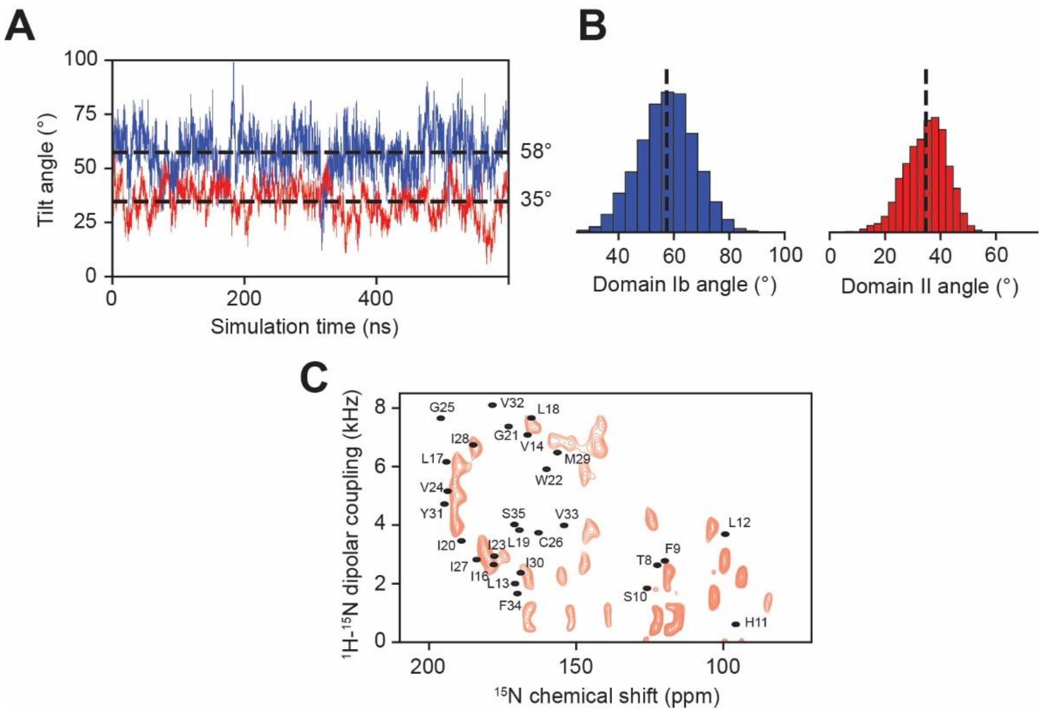


**Figure S4. (A)** Domain Ib (blue) and II (red) tilt angles measured over 600 ns of unrestrained MD simulation and **(B)** corresponding histograms. (C) SLF spectrum back-calculated from unrestrained MD simulation (black) overlaid onto an experimental (SE)-SAMPI4 spectrum of $^{15}$N in flipped bicelles (red).

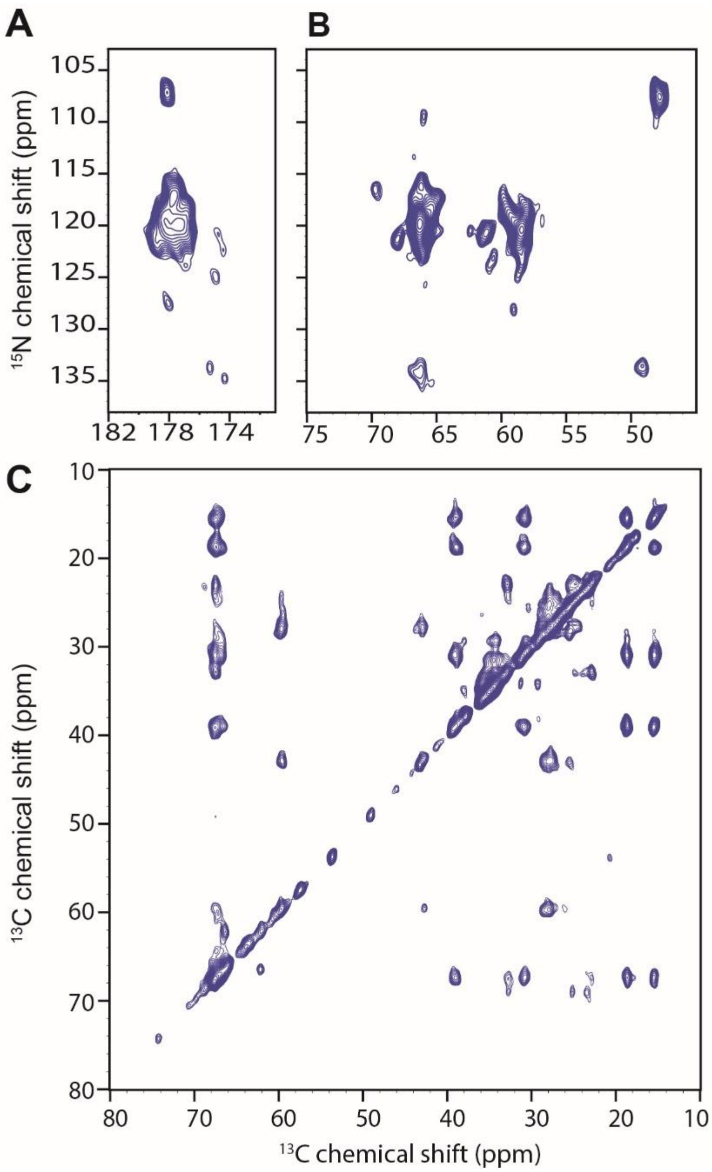


**Figure S5.** MAS-ssNMR spectra of $^{13}C$-$^{15}N$ DWORF reconstituted into $d_{54}$-DMPC liposomes. TEDOR-NCA spectrum showing **(A)** carbonyl and **(B)** Cα regions. **(C)** DARR spectrum with 100 ms mixing time.

**Table S1:** Peak assignments for (SE)-SAMPI4 experiments of DWORF used as restraints in simulated annealing and/or RAOR-MD refinement.

| Residue | DWORF | | DWORF-P15A | |
|---|---|---|---|---|
| | CS | DC | CS | DC |
| G6 | 103.0 | 1.55 | 104.8 | 1.56 |
| S7 | 93.7 | 2.23 | 94.8 | 1.96 |
| T8 | 99.5 | 2.81 | 101.9 | 2.07 |
| F9 | 118.5 | 2.43 | - | - |
| S10 | 115.5 | 1.16 | 114.4 | 1.05 |
| H11 | 85.2 | 1.37 | - | - |
| L12 | 103.9 | 3.86 | - | - |
| L13 | 174.5 | 2.89 | - | - |
| V14 | 150.0 | 6.72 | 143.5 | 5.85 |
| I16 | 166.3 | 1.16 | 158.4 | 0.62 |
| L17 | 189.7 | 6.02 | 177.4 | 5.04 |
| L18 | 142.2 | 6.67 | 128.1 | 5.57 |
| L19 | 147.4 | 2.98 | - | - |
| I20 | 181.7 | 3.17 | 180.4 | 3.26 |
| G21 | 157.3 | 6.89 | 147.0 | 6.43 |
| W22 | 147.3 | 5.83 | 133.0 | 4.52 |
| I23 | 166.9 | 2.28 | 157.5 | 1.12 |
| V24 | 190.4 | 4.96 | 181.5 | 4.15 |
| G25 | 142.2 | 7.31 | 129.1 | 6.73 |
| C26 | 146.4 | 3.88 | 134.3 | 2.59 |
| I27 | 179.3 | 2.73 | 172.7 | 2.01 |
| I28 | 184.4 | 6.69 | 176.0 | 5.89 |
| M29 | 143.0 | 6.97 | 129.7 | 5.98 |
| I30 | 154.7 | 2.22 | 150.3 | 0.79 |
| Y31 | 190.6 | 3.98 | 184.2 | 3.16 |
| V32 | 165.2 | 7.43 | 155.1 | 6.79 |
| V33 | 124.6 | 4.09 | 114.3 | 3.12 |
| F34 | 139.0 | 1.05 | - | - |
